# A multiomic network approach uncovers disease modifying mechanisms of inborn errors of metabolism

**DOI:** 10.1101/2025.02.19.639093

**Authors:** Aaron Bender, Pablo Ranea-Robles, Evan G. Williams, Mina Mirzaian, J. Alexander Heimel, Christiaan N. Levelt, Ronald J. Wanders, Johannes M. Aerts, Jun Zhu, Johan Auwerx, Sander M. Houten, Carmen A. Argmann

## Abstract

For many inborn errors of metabolism (IEM) the understanding of disease mechanisms remains limited in part explaining their unmet medical needs. We hypothesize that the expressivity of IEM disease phenotypes is affected by the activity of specific modifier pathways, which is controlled by rare and common polygenic variation. To identify these modulating pathways, we used RNA sequencing to generate molecular signatures of IEM in disease relevant tissues. We then integrated these disease signatures with multiomic data and gene regulatory networks generated from animal and human populations without overt IEM. We identified and subsequently validated glucocorticoid signaling as a candidate modifier of mitochondrial fatty acid oxidation disorders, and we re-capitulated complement signaling as a modifier of inflammation in Gaucher disease. Our work describes a novel approach that can overcome the rare disease-rare data dilemma, and reveal new IEM pathophysiology and potential drug targets using multiomics data in seemingly healthy populations.

Take-home message: Multiomic data and gene regulatory networks derived from animal and human populations without overt IEM can be used to study disease mechanisms in inborn errors of metabolism.

## Introduction

Inborn errors of metabolism (IEM) are a collection of rare genetic disorders involving biochemical processes. IEM have historically been considered single-gene diseases with “simple” Mendelian, most often autosomal recessive, inheritance patterns. As such the severity of the clinical presentation is assumed to correlate with mutation deleteriousness also known as genotype-phenotype correlation.^1–4^ Research on IEM has therefore primarily focused on the known defective enzyme or pathway. Although this has been a highly effective strategy, the disease mechanisms underlying the clinical presentation of IEM often remain ill-defined. In addition, IEM present as a spectrum of disease phenotypes ranging from severe to mild or attenuated, sometimes without a distinct genotype-phenotype correlation. Consequentially, treatment is often suboptimal, addressing the symptoms rather than the underlying disease. The unexplained variation in genotype-phenotype correlation illustrates that factors (e.g. genetic or environmental) other than the defective enzyme or pathway may influence the disease course of the IEM. Accordingly, diet and feeding status are well-established disease modifiers in many IEM.

Genome-wide association studies (GWAS) have demonstrated that common genetic variants affect metabolite concentrations and as such determine an individual’s metabolic individuality.^5,6^ These variants are often identified in the same genes in which rare variants cause IEM with the same metabolite perturbations, although with differing effect sizes.^5,6^ Similarly, many disease-associated recessive variants can produce mitigated phenotypes in heterozygous carriers.^7–9^ In addition, susceptibility loci underlying common disease were found to be globally enriched in Mendelian disease loci.^10,11^ Thus, variation in metabolic genes exists in allelic series as part of a continuum of biochemical phenotypes with common variation on one extreme associated with subtle phenotypes and rare variants on the other extreme giving rise to IEM.^1^ As such IEM are comparable to complex diseases, as they are emergent phenotypes driven by a primary disease gene and influenced by multiple (genetic or other) modifiers.^1–4^ Re-categorization of IEM as part of a spectrum of metabolic phenotypes alongside common metabolic diseases motivates studying IEM like common disease, with data-driven, multiomic approaches.^12–23^ In this way the search space for IEM relevant pathophysiology can be guided beyond the known defective enzyme. We can also overcome the rare disease-rare data hurdle^24^ as common disease experimental model systems and their associated multiomic datasets can be repurposed to inform on novel IEM modifying biology.^1^

The mouse is a rich resource of multiomic datasets that can be used to study complex traits. One example is the BXD recombinant inbred (RI) genetic reference population of mice, which consists of nearly 200 strains generated from C57BL/6J (B6) and DBA/2J (D2) (here referred to as BXD) and encompasses ∼5 million SNPs.^25,26^ Available multiomic data include genotype, multiple tissue transcriptome and proteome data, and clinical phenotyping measured in animals under different dietary challenges.^27–29^ For this study, we have measured the abundance of ∼60 different metabolites in plasma samples of 40 BXD RI strains. These metabolites include acylcarnitines, amino acids, and glycosphingolipids, which are considered diagnostic biomarkers for IEM such as mitochondrial fatty acid oxidation (FAO) disorders, organic acidemias, aminoacidopathies and some lysosomal storage disorders. The BXD multiomic data were then integrated with IEM disease signatures to reveal novel modifying biology.

## Materials and Methods

Detailed materials and methods are available in the supplementary material.

### BXD RI strains

The BXD RI cohort was described previously.^27,28^ Plasma was pooled equally by volume for all animals to create sufficient quantities for all metabolite analyses.

### Metabolite measurements

Sixty-five plasma metabolites and 12 liver metabolites were measured in the BXD RI cohort as described in the supplementary material. All data are available on GeneNetwork.org under the “BXD Published Phenotypes” set.

### QTL assessment in the BXD RI cohorts

Metabolite measurements were correlated with genotype for single nucleotide polymorphisms (SNPs) at 7,320 molecular markers in linkage disequilibrium across the genome using the Haley-Knott regression model of normally distributed traits using the software package R/qtl.^30^

### RNAseq of C57BL/6J and DBA/2J liver and LCAD KO liver and gastrocnemius muscle

Total RNA from liver and gastrocnemius muscle was isolated using QIAzol lysis reagent followed by purification using the RNeasy kit (Qiagen). RNA was submitted to the Genomics Core Facility at the Icahn School of Medicine at Mount Sinai for further processing. mRNA-focused cDNA libraries were generated using Illumina reagents (polyA capture), and samples were run on an Illumina HiSeq 2500 sequencer to yield appropriate read depth. All sequencing data are available in the GEO database (GSE186973, GSE186613 and GSE186648).

### Long read sequencing

The sequence of the retrotransposon in *Mlycd* was determined using PacBio Single Molecule, Real-Time (SMRT) sequencing and is available at GenBank (MH036232).

### Generation and characterization of a congenic DBA/2J line with Mlycd^B6/B6^

In the BXD6 strain, the C57BL/6J marker allele gnf08.119.598 was isolated in a DBA/2-context using speed congenics.

### Generation of IEM disease signatures

The LCAD KO signatures were derived from a comparison of WT with LCAD KO mice. Significant DEGs were defined using an adjusted p value < 0.05 (Benjamini-Hochberg method) with no fold change cut-off.

### Generation of coexpression networks from the BXD cohort

Plasma metabolites and liver and muscle gene expression were organized into modules using WGCNA.^31^

### Bayesian gene regulatory networks (GRNs)

Mouse Bayesian GRNs were generated as previously described from the liver and muscle gene expression data generated from a series of segregating mouse populations.^32^ The human liver GRN was constructed from the human liver cohort comprised of 427 Caucasian subjects.^32^ The muscle GRN was newly constructed as described^33^ from muscle gene expression data obtained from the publicly available datasets on the Genotype-Tissue Expression (GTEx) portal.

### Generation of tissue-specific, disease-specific subnetworks

Liver and muscle BXD coexpression modules were tested for enrichment in IEM disease signatures using a one-side Fisher’s exact test. P values were corrected for multiple testing using the Benjamini-Hochberg method.

### Shortest path analysis

Shortest path analysis using the distances function of the R package igraph was used to evaluate how well GRNs captured IEM disease signatures and associated gene coexpression modules.

### Annotation of gene sets, coexpression modules and subnetworks

Gene sets, coexpression modules and subnetworks were annotated for function using enrichment analysis and several pathway databases as described in the supplementary material.

### Key driver analysis (KDA)

Key driver analysis was performed using the KDA library in R.^34^

### Transcription factor binding motif enrichment: iRegulon

Transcription factor binding motif enrichment was performed using ChIP-seq derived gene sets in iRegulon.^35^

### Validation of the role of glucocorticoid signaling during food withdrawal in a mouse model with a FAO defect

Mice (5 males and 2 females per group) were treated with L-aminocarnitine and/or mifepristone as described in the supplementary material.

## Results

### Variation in the *Mlycd* controls malonylcarnitine levels in BXD RI strains

The most intuitive and direct source of potential IEM modifiers are genes that control levels of IEM-relevant metabolites. We have previously observed many naturally occurring biochemical traits in inbred mouse strains including 2-aminoadipic and 2-oxoadipic aciduria due to *Dhtkd1* deficiency.^36^ We measured amino acids, acylcarnitines, and (glyco)sphingolipids in plasma samples of 40 different BXD strains on chow and high fat diet (CD and HFD^27,28^, **Table S1**). In liver samples from the same cohort, we measured (glyco)sphingolipids as well as the activity of β-hexosaminidase (HEX) and β-glucocerebrosidase (GBA), which are both lysosomal enzymes involved in glycosphingolipid degradation. Animals on two different diets (CD and HFD) were used to maximize the range of variation in metabolite levels across the cohort as environmental exposures such as diet are known to have a profound effect on metabolite abundance. Indeed, hierarchical clustering analysis illustrated that diet had a primary influence on metabolite abundance compared to the genetic diversity across the strains (**Figure S1A**). The genetic control of these metabolites can be studies through quantitative trait locus (mQTL) mapping (**Table S2, S3**). In order to identify candidate genes that may control the abundance of the metabolites, additional approaches are often necessary. These include the use of gene expression as a quantitative trait (eQTL mapping, **Table S4-7**), co-mapping of mQTLs and eQTLs (**Table S8**), correlation between gene expression and metabolite abundance (**Table S9**) as well as canonical knowledge of the function of genes within the associated genomic region.

We observed a novel diet-independent mQTL for plasma C3DC (malonylcarnitine) mapping to a region of chromosome 8 (**Figure 1A, Table S2, S3**) previously associated with liver levels of malonate and methylmalonate.^37^ This region contains *Mlycd* as the sole candidate causal gene. *Mlycd* encodes malonyl-CoA decarboxylase (MCD), which is an enzyme that catalyzes the conversion of malonyl-CoA into acetyl-CoA. Elevated plasma C3DC and increased urine malonate and methylmalonate are classic markers of human MCD deficiency.^38^ Variation in *Mlycd* expression is the mechanism underlying the genetic control of C3DC levels in the BXD RI mouse population as established by a co-localizing liver *cis* eQTL for *Mlycd* (**Table S4, S5**) and a significant negative correlation between *Mlycd* expression and C3DC levels (**Figure 1B, S1B, Tables S8, S9**). Haplotype mapping at this physical location shows that high plasma C3DC levels segregate with the presence of the parental DBA/2J allele in a Mendelian fashion (**Figure 1B, S1B**). Accordingly, malonate was higher in urine samples from DBA/2J mice when compared to urine samples from C57BL/6J (**Figure S1C**). Interestingly, the *cis* eQTL for *Mlycd* is tissue specific and observed in liver, lung and spleen, but not in other major metabolic organs such as muscle and heart (for muscle see **Table S6, S7**).

**Figure 1.**
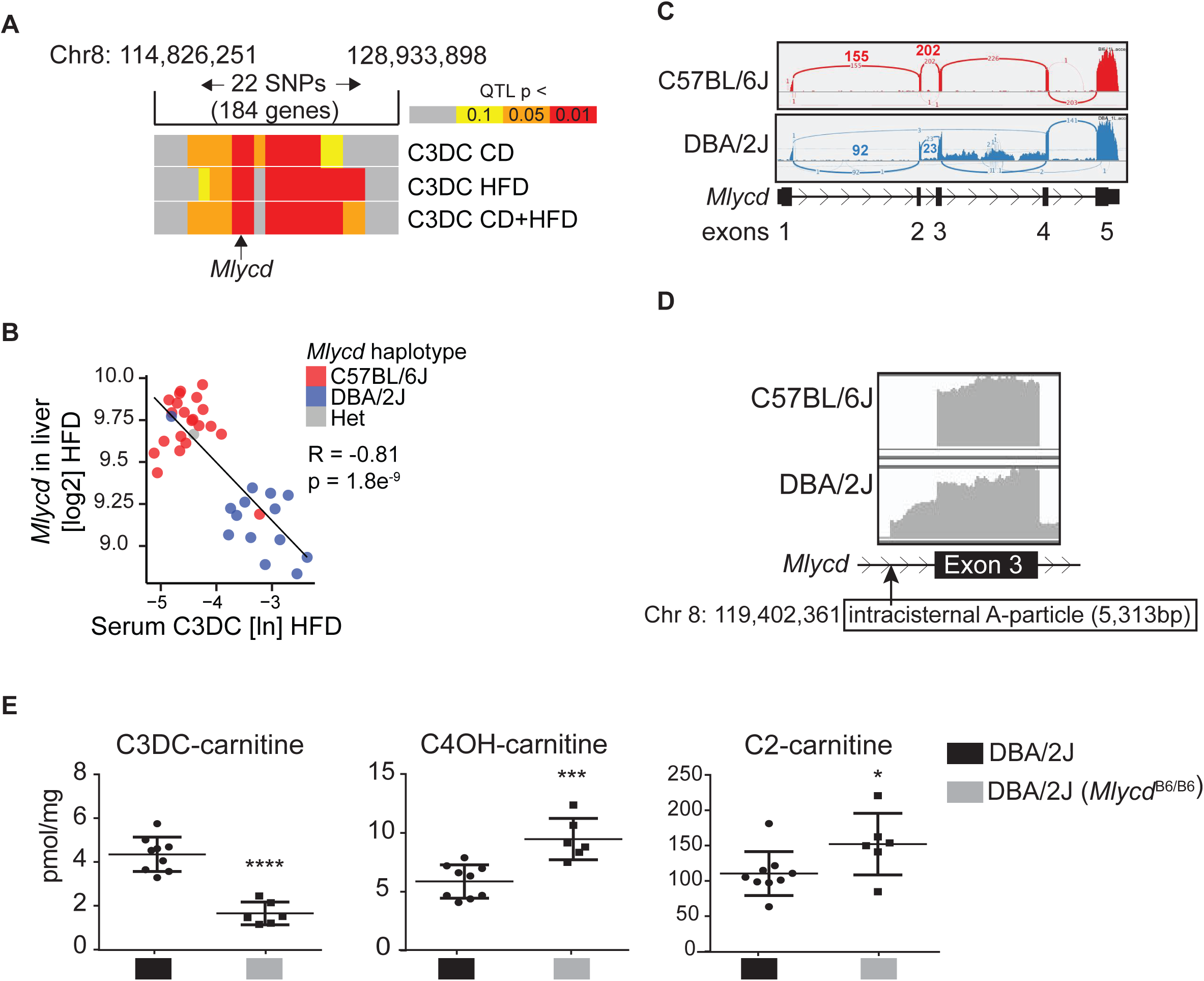
Identifying potential modifiers of IEM metabolite abundance through QTL mapping. (A) QTL mapping for C3DC on a region of chromosome 8 containing *Mlycd*. (B) The significant negative correlation between plasma C3DC levels and *Mlycd* gene expression in the liver of BXD RI mice on HFD. The animals almost perfectly separated expression levels according to haplotype near the *Mlycd* gene. Animals with the DBA/2J haplotype had low *Mlycd* expression and high C3DC, whereas the opposite relationship was observed for C57BL/6J. (C) A Sashimi plot visualizing the splice junctions and read density from aligned RNA-seq data of liver tissue from a C57BL/6J and a DBA/2J mouse. All exons and introns from the *Mlycd* gene are displayed. (D) A detail of the Sashimi plot showing read density in the region around *Mlycd* exon 3. Many intronic reads are observed in the DBA/2J liver sample. The position of the identified insertion of an intracisternal A-particle in intron 2 of *Mlycd* in the DBA/2J genome is indicated. (E) Liver abundance of C3DC, C4OH and C2-carnitine in DBA/2J mice and mutant DBA/2J mice carrying the C57BL/6J allele for *Mlycd*.

In order to identify the variant that causes MCD deficiency in the BXD strains, we used RNA sequencing data generated from liver of the parental C57BL/6J and DBA/2J strains. We compared reads that mapped to the region containing *Mlycd*^36^ and observed more reads in introns 2 and 3 in DBA/2J mice when compared to C57BL/6J (**Figure 1C, 1D**). Using PCR on genomic DNA, we found a 5,313 bp insertion in intron 2 of *Mlycd* from DBA/2J mice (chr8:119,402,361), which was not present in C57BL/6J or 129S2/SvPasCrl mice (**Figure S1D**). We obtained the full sequence of this insert using long-read sequencing, which revealed an intracisternal A-particle (IAP) retrotransposon with identical long-terminal repeats and a 3,204 bp open reading frame encoding the gag-pol fusion protein (**Figure 1D**, GenBank accession number MH036232).^39,40^ Elevated C3DC was reported previously in SM/J mice.^41^ We detected a similar structural variant in genomic DNA from SM/J mice (**Figure S1D**), which likely reflects a close phylogenetic relationship with DBA/2J.^42^

To further validate the link between the DBA/2J variant of *Mlycd* and plasma levels of C3DC, we compared hepatic levels of acylcarnitines in wild-type (WT) DBA/2J mice with levels in congenic DBA/2J mice carrying the C57BL/6J *Mlycd* allele (*Mlycd*^B6/B6^) after overnight food withdrawal (**Figure 1E**). DBA/2J mice with *Mlycd*^B6/B6^ had lower levels of C3DC when compared to WT DBA/2J mice, likely as a result of restored MCD activity (**Figure 1E**). The levels of hydroxybutyrylcarnitine (C4OH-carnitine) and acetylcarnitine (C2-carnitine) were higher in DBA/2J mice with *Mlycd*^B6/B6^ when compared to WT DBA/2J mice (**Figure 1E**). In the BXD RI population on CD, liver *Mlycd* expression significantly correlated with C4OH-carnitine as well (*r* = 0.51, **Figure S1E**). The increased C4OH-carnitine and C2-carnitine likely indicate higher FAO as these acylcarnitines represent ketone bodies and acetyl-CoA, which are both FAO end-products.^43^ These results are consistent with MCD activity controlling mitochondrial FAO and ketogenesis in the liver through modulation of malonyl-CoA levels and, as a consequence, the activity of the rate-limiting hepatic carnitine palmitoyltransferase 1 (CPT1A) (**Figure S1F**).^44,45^ As such, functional variants in *Mlycd* could be potential modifiers of clinical phenotypes of FAO disorders, in particular fasting-induced hypoketosis.

### Unbiased identification of IEM modifying biology through integration of multiomics data

The identification of variants controlling IEM-related metabolites is one source potential modifying biology. A potential disadvantage of this approach is that the selection of these metabolites is biased because it is based on existing knowledge of disease biology. To overcome this limitation, we next interrogated these population-based multiomics data with unbiased IEM disease signatures. To generate a novel disease signature for a FAO disorder, we used an animal model. We and others have shown that the long-chain acyl-CoA dehydrogenase (LCAD or *Acadl*) knockout (KO) mouse presents with phenotypes that resemble specific aspects of FAO disorders such as very long-chain acyl-CoA dehydrogenase (VLCAD) deficiency. These phenotypes include increased long-chain acylcarnitines, fatty liver, fasting-induced hypoketotic hypoglycemia and cardiac hypertrophy.^46–49^ Following overnight food withdrawal to increase dependence on FAO, LCAD KO mice and WT controls were euthanized and organs were collected.^50^ To generate a molecular disease signature underlying the modeled IEM, we performed RNA-sequencing analysis on liver and gastrocnemius muscle, which are both organs that play a prominent role in the pathophysiology of long-chain FAO disorders. In liver, we identified 2,633 significantly differentially expressed genes (DEGs), of which 1,355 were up- and 1,278 were down-regulated (**Table S10**). The DEG signature confirms previously reported changes in hepatic glucose metabolism of LCAD KO mice such as decreased expression of *Gck*, *Slc2a2*, *Pklr* and *Pygl*, and increased expression of *Pdk4*.^48^ In muscle, we identified 327 significantly DEGs, of which 207 were up- and 120 were down-regulated (**Table S10**). Among the top upregulated genes were *Acot1* and *Acot2*, which encode acyl-CoA thioesterases involved in fatty acid metabolism. These DEG signatures reflect the molecular changes in liver and muscle of the fasted LCAD KO mouse.

We next focused on identifying potential FAO disease modifying biology in the multiomics BXD RI data. For this, we first organized the plasma metabolite and liver and muscle gene expression data from the BXD RI strains through weighted correlation network analysis (WGCNA^31^). The WGCNA algorithm yields a network of different modules consisting of highly correlated metabolites or transcripts. Because of this, the metabolites or transcripts within a module are likely coregulated and therefore functionally related. The metabolite network was built from combined CD and HFD cohorts and revealed five distinct modules (**Table S11**, and **Figure S2**), which were named according to their main constituents; long-chain acylcarnitines (LCAC, turquoise), short-chain acylcarnitines (SCAC, blue), lipids (yellow), branched-chain amino acids (BCAA, green), and total amino acids (total AA, brown). Each module’s eigenvector (first principal component) explained, on average, 62 to 76% of the variability. Two metabolite modules strongly responded to diet, the lipids and SCAC modules (**Figure S2**). Liver and muscle gene co-expression networks were generated for the CD and HFD cohorts separately (**Table S12**). In general, the modules were found conserved between diets implicating diet-independent gene coordination across a wide array of pathways and functions (**Figure S3, Table S13**). Expression values for all genes within a module were collapsed into a single value eigenvector for each BXD RI strain (**Tables S14-S17**) and associated module QTLs calculated (**Table S18**).

We then determined which gene co-expression modules were significantly enriched in the LCAD KO liver and muscle DEGs. The most significantly enriched modules in liver were module 3 in CD (LC3) and module 2 in HFD (LH2). Module 4 in CD (MC4) and module 2 in HFD (MH2) were most significantly enriched in muscle (**Figure 2A, 2B**, **Table S19**). Gene composition of these liver and muscle modules was conserved (**Figure S3, Table S13**). Further investigation of these modules revealed they were significantly correlated to the LCAC and total AA metabolite modules (**Figure 2A, 2B**, **Table S20**) and to the clinical trait fasting-induced weight loss (FIWL).

**Figure 2.**
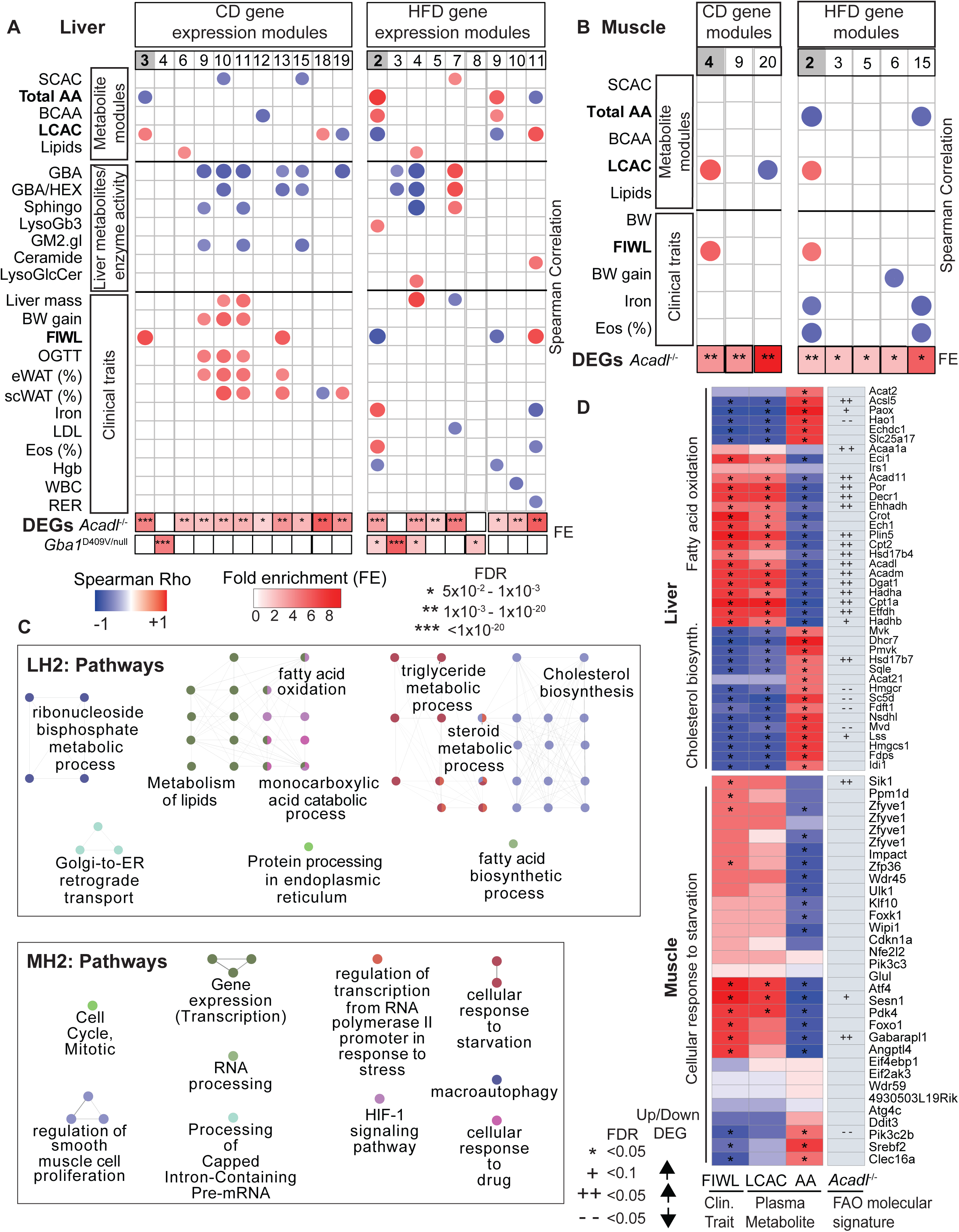
Identifying IEM disease modifying biology through co-mapping IEM relevant metabolites, traits and molecular signatures to BXD gene coexpression modules. (A) Gene coexpression modules from BXD liver were tested for enrichment in LCAD KO and Gaucher disease DEGs (color according to fold enrichment). Gene module eigengenes (PC1 calculated for each module) were correlated to metabolite module eigenmetabolites (PC1 for each module), liver metabolite and enzyme activity levels, and clinical traits associated with different IEM (circle color according to Spearman’s r correlation; circle size is proportionate to FDR significance). (B) Gene coexpression modules from BXD muscle were tested for enrichment in DEGs from LCAD KO muscle (color according to fold enrichment). Gene module eigengenes were correlated to metabolite module eigenmetabolites and clinical traits associated with different IEM (circle color according to Spearman’s r correlation; circle size is proportionate to FDR significance). (C) LCAD KO-specific coexpression modules liver HFD 2 (LH2) and muscle HFD module 2 (MH2) were annotated with enriched GO, KEGG, and Reactome gene sets. Displayed are clusters of pathways that have similar gene membership. (D) Expression levels of genes from LH2 that map to “fatty acid oxidation” and “cholesterol biosynthesis” gene sets and genes from MH2 that map to the gene set for “cellular response to starvation” were correlated to fasting-induced weight loss (FIWL), the long-chain acylcarnitine (LCAC) metabolite module, and the total amino acid (AA) metabolite module assuming non-normal distribution of gene expression (Spearman’s r). P values were corrected for multiple testing using the Benjamini-Hochberg method. Each gene was annotated for membership in the LCAD KO disease signatures.

Next the gene expression modules were functionally annotated through pathway enrichment analysis. In the liver, the LH2 module, which shares many genes with LC3, was enriched for genes functioning in “fatty acid oxidation”, “cholesterol biosynthesis”, “protein processing in endoplasmic reticulum”, amongst others (**Figure 2C upper panel, Table S21**). LH2 genes belonging to the FAO pathway, such as liver *Acadl* and *Cpt2* were positively correlated with LCAC and FIWL and negatively correlated with total AA. The exact opposite pattern was observed for *Hmgcr* and *Dhcr7*, which are genes involved in the cholesterol biosynthesis pathway (**Figure 2D**, **Table S21**). In the muscle, the MH2 module, which shares many genes with MC4, was enriched for metabolic functions such as “cellular response to starvation”, “macroautophagy” and “HIF-1 signaling pathway” (**Figure 2C lower panel, Table S21**). MH2 genes associated with autophagy such as *Foxo1*, *Sesn1* and *Ulk1*, were positively correlated with FIWL and LCAC and negatively with total AA (**Figure 2D**). Notably, the total AA and LCAC modules were anti-correlated indicating that mice with relatively high levels of AAs in general have lower levels of LCAC (**Figure 2D**). In summary, the modules enriched in the LCAD KO disease signatures highlight genes implicated in the coordinated physiologic response of liver and muscle to starvation. During starvation, hepatic FAO is induced in order to enable ketogenesis, whereas cholesterol biosynthesis is repressed. Muscle proteolysis enabled by autophagy is responsible for the generation of gluconeogenic AAs. Importantly, our analysis uncovers that in the BXD RI mouse cohort there is inherent population-level variation in the response to starvation. Because fasting is a critical environmental trigger for pathology in FAO disorders, we hypothesize that we can leverage the highlighted muscle and liver gene expression modules and associated processes to identify important modifying biology underlying FAO disorders.

### Bayesian gene regulatory networks reveal glucocorticoid signaling as a putative modifier of FAO disorder biology in the muscle

To prioritize the genes/pathways amongst the muscle DEGs and modules associated with FAO disorders, we utilized Bayesian gene regulatory networks (GRNs). GRNs are mathematical models often represented as network graphs that predict the regulatory interaction between genes. GRNs can incorporate genetic information such as eQTLs as causal anchors during network construction in order to statistically infer directionality in gene pair relationships. The GRNs we surveyed were generated from muscle expression data generated in second generation (F2) offspring of several mouse inbred strains or from the human GTEx project.^32,51^ We first projected the LCAD KO muscle DEGs as well as the genes of the MC4 and MH2 modules onto the muscle GRNs and calculated the average shortest path length between all possible gene pairs and compared that to the average shortest path length of random sets of genes of similar size. We found the mean shortest path length between any pair of nodes to be less for the DEGs and modules than for randomly sampled gene sets. This was significant (z < - 2) for all 3 gene sets in the mouse GRN, and also for 1 gene set in the human GRN (**Figure 3A, S4A, Table S22**). Shorter path lengths between genes in a network context reflect that the nodes are more connected and thus more likely to be coregulated. Importantly, this observation in both mouse and human muscle GRNs, is a confirmation that the gene expression coordination in response to a severe single gene perturbation (LCAD KO) can be recapitulated in networks constructed from individuals without an overt mitochondrial FAO defect. Critically, this finding supports the further use of these GRNs to study the molecular response to a genetic mitochondrial FAO defect.

**Figure 3.**
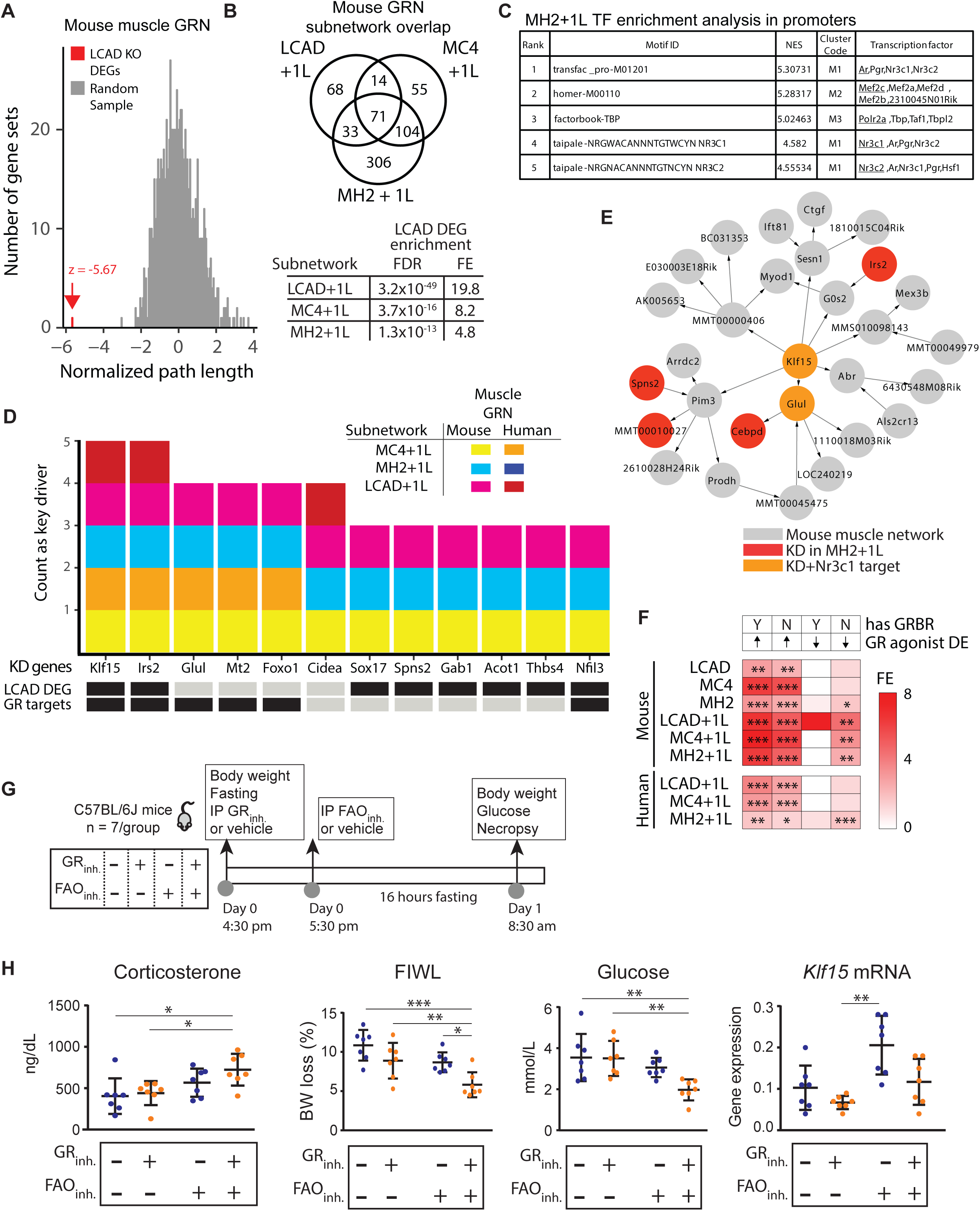
Bayesian gene regulatory network analysis reveals glucocorticoid signaling as a candidate modifying pathway in FAO-deficient muscle. (A) The normalized path length of LCAD KO muscle DEGs is significantly smaller than the normalized path lengths of random gene sets of equal size in the mouse muscle GRN. (B) Mouse subnetworks generated from LCAD KO muscle DEGs and the MC4 and MH2 modules share many genes (top) and are highly enriched in the original LCAD KO muscle DEGs (bottom; FE = fold enrichment). (C) Genes from MH2 + 1L subnetwork generated from the mouse muscle GRN were tested for enrichment in regulatory motifs using the iRegulon feature of Cytoscape, and the top five most enriched motifs are shown (NES = normalized enrichment score). (D) The top ranked key drivers (KD) of LCAD KO subnetworks are shown according to the frequency of identifying the gene as a key driver across the 6 subnetworks. Key drivers are annotated for being a member of the LCAD KO muscle DEGs and for being a GR target gene. (E) The local network structure from the mouse muscle GRN around the top-ranked key driver *Klf15* reveals multiple gene targets of glucocorticoid receptor such as *Glul*. (F) LCAD KO muscle subnetworks were tested for enrichment in GR responsive genes (FE = fold enrichment). The curated GR-responsive gene set was separated into genes upregulated (arrow up) or downregulated (arrow down) following treatment with a GR agonist and genes that contained a glucocorticoid receptor binding region (GRBR) (Y) or not (N) as determined by ChIP-seq. (G) Schematic representation of the mouse experiment that validates the role of GR as a modifier of the fasting response in animals with a FAO deficiency. At 4:30pm, mice were treated with the GR inhibitor mifepristone (GR_inh._) or vehicle. One hour later (5:30pm) mice received an intraperitoneal (ip) injection of L-AC (FAO_inh_.) or vehicle. Mice were euthanized after overnight food withdrawal. (H) Plasma corticosterone FIWL, plasma glucose, and muscle *Klf15* liver mRNA levels were measured in the mice treated as described above. The ANOVA F value was significant for all 4 traits (Table S31). Indicated are the results for the Tukey’s multiple comparisons test.

By projecting the LCAD KO molecular disease signatures onto GRNs and including nearest neighboring genes that are not part of the signature, we can study the molecular response to a long-chain FAO defect in the form of a disease-specific subnetwork and also increase the search space for modifying biology. The LCAD KO muscle DEG signature as well as the MC4 and MH2 coexpression modules were projected onto both mouse and human muscle GRNs, and subnetworks were extracted including all seed set genes and their nearest neighbors (i.e. + one layer (1L), **Table S23**). Consistent with our shortest path analysis, the mouse and human subnetworks of the LCAD KO DEGs+1L, MC4+1L and MH2+1L were significantly enriched in LCAD KO muscle DEGs (**Figure 3B, S4B, Tables S24, S25**).

To help prioritize the biology underlying the various LCAD KO-associated subnetworks, we characterized the genes according to their enrichment in canonical transcription factor-binding motifs. A motif predicted to bind the glucocorticoid receptor (GR or nuclear receptor subfamily 3 group C member 1, Nr3c1) belonged to the most enriched cluster of motifs (M1) in the MC4+1L and MH2+1L subnetworks and the eighth most enriched cluster (M8) from the LCAD KO DEGs+1L subnetwork (**Figure 3C, Table S26**). Using the structure of the GRNs, we then identified key driver genes, defined as those genes which are the most highly connected to the genes of the disease subnetworks. The hypothesis underlying key driver analysis is that disruption of key driver genes would have a greater effect on the phenotypes captured by the disease subnetworks than non-key driver genes.^34^ Key driver analysis was performed for the disease-associated subnetworks LCAD KO DEGs+1L, MC4+1L and MH2+1L on both the mouse and human muscle GRNs. Key drivers were ranked according to the number of times a gene was identified. They were also annotated for membership of the LCAD KO signature and a curated set of GR target genes^52^ (**Figure 3D, Table S27, S28**). Krüppel-like factor 15 (*Klf15*) was identified as a key driver in 5 of 6 subnetworks. *Klf15* was differentially upregulated in the LCAD KO muscle, is a predicted target of GR, and is known to mediate the response to starvation in the muscle.^53–55^ The network neighbors of *Klf15* in the mouse muscle GRN include *Glul* and *Irs2* (**Figure 3E**), which are also key drivers and GR target genes. A more directed survey revealed that LCAD KO-associated signatures, modules, and subnetworks were enriched in genes up-regulated following GR agonist stimulation including a subset of genes containing GR binding domains (**Figure 3F, Table S29**).

If *Klf15* is a key driver of FAO-associated biology, then FAO-associated subnetworks should be enriched in genes that are differentially expressed when *Klf15* expression is perturbed.^56^ Indeed, LCAD KO DEGs, MC4 and MH2 modules, and LCAD KO DEGs+1L, MC4+1L, and MH2+1L subnetworks were enriched in DEGs from skeletal muscle of a *Klf15^−/−^*mouse (**Table S30**). Overall these data implicate GR and KLF15 signaling as potential key modulators of these LCAD KO-associated subnetworks in the muscle. This suggests that modulation of GR signaling could have a significant effect on the phenotype of the LCAD KO mice.

### *In vivo* validation of GR signaling as a modifier of a mouse model with a FAO defect

We next tested the hypothesis that the GR signaling pathway plays a role in the gene expression changes and phenotypes of the LCAD KO mouse model, and therefore may represent modifying biology in long-chain FAO defects. We treated WT mice with the GR antagonist mifepristone (GR_inh_) and/or the mitochondrial FAO inhibitor L-aminocarnitine^57–60^ (FAO_inh_) followed by overnight food withdrawal to induce a catabolic stress condition (**Figure 3G**). Analysis of variance (ANOVA) for each measurement (corticosterone, FIWL, plasma glucose, and *Klf15* mRNA) indicated significant differences between groups (**Table S31**). Following overnight food withdrawal and FAO inhibition, mice showed increased plasma corticosterone levels consistent with induced glucocorticoid signaling to maintain euglycemia (**Figure 3H**). Mice treated with the combination of FAO_inh_ and GR_inh_ lost significantly less weight and were significantly more hypoglycemic, supporting the role of GR signaling to maintain euglycemia through the generation of free AAs through protein degradation and subsequent gluconeogenesis. *Klf15* mRNA expression was increased by inhibition of FAO, which is consistent with its induction in LCAD KO muscle. The decrease in *Klf15* expression upon GR inhibition is consistent with *Klf15* being a GR target gene. Taken together, this animal experiment supports the role of GR signaling as a modifier of the phenotype of mitochondrial FAO disorders through modulating muscle protein catabolism.

### Unbiased identification of disease modifiers in Gaucher disease

To illustrate that our approach is generalizable and applicable to other IEM, we highlight its successful application to recapitulate disease-modifying pathways of Gaucher disease (GD), one of the most common lysosomal storage disorders. We used the 840 DEGs (627 up- and 213 down-regulated) obtained from the liver of *Gba1* p.D409V/null mice as a GD signature^61^ (**Table S32**). Five BXD liver modules, LC4, LH2, LH3, LH4 and LH8 were enriched for GD DEGs (**Figure 2A, Table S19**). Whereas LH8 and LH3 were conserved in chow diet as LC4 (**Figure S3, Table S12**), the genes belonging to LH4 were dispersed over multiple chow modules (LC3, LC9, LC10 and LC11, **Figure S3**). Modules LH3 and LH4 significantly correlated to the activity of GD-deficient enzyme GBA (**Figure 2A, Table S20**). In addition, the LH4 module correlated to sphingolipids and lysoglucosylceramide and, importantly, to liver mass, a proxy measurement for hepatosteatosis. Hepatosplenomegaly amongst other patterns of liver damage (hepatocarcinoma and fibrosis) are common, heterogeneous phenotypes of GD patients. LH4 was also correlated to the plasma lipid module (**Figure 2A, Table S20**), representing a connection between hepatic lipid metabolism and plasma metabolite levels capturing the molecular and phenotypic characteristics of GD.

GD DEGs and associated BXD modules are tightly co-regulated in liver GRNs generated from population models of mouse^62^ and human^63^ as demonstrated using shortest path analysis (**Figure 4A, Figure S5, Table S33**). They were also used to isolate GD-associated subnetworks through the projection of GD DEGs or associated modules onto GRNs and including nearest neighbors (**Table S34**). The resulting GD-specific, tissue-specific subnetworks share many genes (**Figure 4B, Table S35**) and were enriched in the GD DEG signature, reflecting that they continued to reflect the IEM disease signature (**Figure 4B**, **Table S36**).

**Figure 4.**
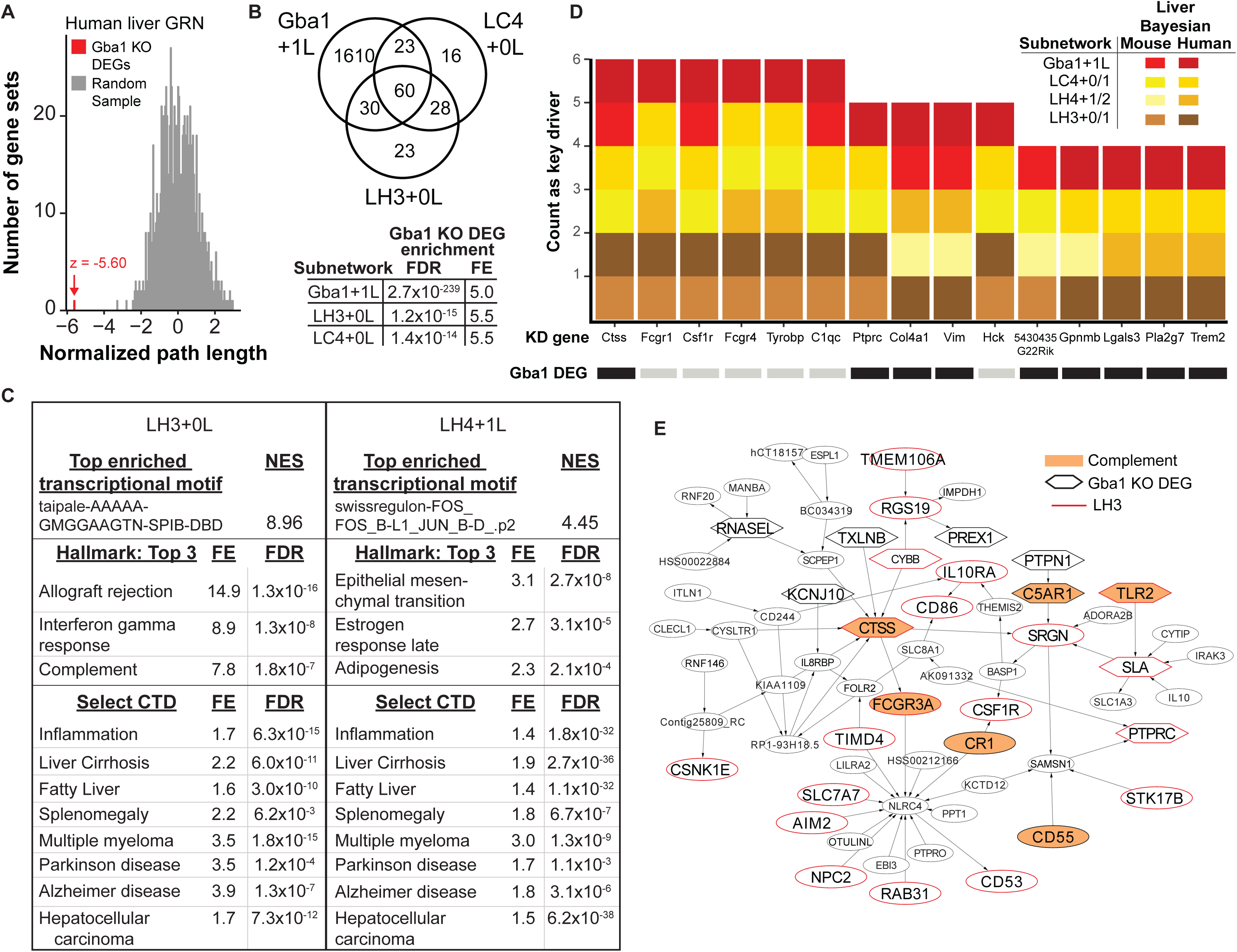
Identifying inflammatory pathways associated with phenotype variability in Gaucher disease. (A) The normalized path length of GD liver DEGs is significantly shorter than the normalized path lengths of random gene sets of equal size in the human liver GRN. (B) Subnetworks generated from integration of the mouse liver GRN with GD liver DEGs and BXD liver coexpression modules enriched in GD liver DEGs share genes in common (top) and are highly enriched in the original GD signature (bottom; FE = fold enrichment). (C) Subnetworks generated from integration of the mouse liver GRN with liver HFD coexpression module 3 (LH3) and liver HFD coexpression module 4 (LH4) were annotated for enrichment in transcriptional motifs, Hallmark gene sets from MSigDB, and pathology-associated gene sets from the Comparative Toxicogenomics Database (CTD). (D) Top-ranked key drivers are shown according to the frequency that they were identified as a key driver across liver-specific, GD-specific subnetworks generated in the human and mouse liver GRNs. Key drivers were annotated for membership in the GD liver DEGs. (E) The local network structure around the top-ranked key driver *CTSS* in the human liver GRN reveals multiple genes belonging to the LH3 subnetwork, the complement gene set from Hallmark, and the GD liver DEGs.

Subnetworks associated with LH3 and GD DEGs were enriched in WikiPathways gene sets “Macrophage markers” and “Microglia Pathogen Phagocytosis Pathway” (**Table S37**) and Hallmark gene sets “allograft rejection”, “interferon gamma response”, and “complement” (**Figure 4C**, **Table S38**). The top enriched transcriptional motif for the LH3+0L subnetwork was a canonical SPIB binding domain with predicted transcription factors including the macrophage development and iron homeostasis regulator Spi-C (SPIC, **Figure 4C, Table S39**). These results are consistent with dysfunctional macrophage activity due to lysosomal accumulation of glucocerebroside.^64^ The LH4+1L subnetwork was most enriched for the pathway terms “epithelial-mesenchymal transition”, “adipogenesis”, “Nuclear receptors in lipid metabolism and toxicity”, and the enriched transcriptional motif was predicted to bind to AP-1 (a variable dimer of Fos and Jun subunits)(**Figure 4C, Table S37, S38**). Epithelial-mesenchymal transition in the liver may be related to chronic inflammation and the development of liver fibrosis. Patients with GD have a significantly greater risk for developing certain cancers (including multiple myeloma and hepatocellular carcinoma^65^) and Parkinson disease.^66^ Interestingly, both the LH3+0L and LH4+1L subnetworks were enriched in genes associated with these diseases (**Figure 4C**, **Table S40**).

Cathepsin S (*Ctss*) was identified as a key driver gene in six of eight total subnetworks tested (**Figure 4D, Table S41**). *Ctss* is upregulated in liver and spleen of the GD mouse model, and serum CTSS levels have been proposed as a biomarker for GD disease severity due to their elevated levels in patients before treatment initiation.^67^ *CTSS* is a target of the SPIC transcription factor, and the *CTSS* subnetwork in human liver GRN (including complement-associated genes included *FCGR3A*, *CR1*, *CD33*, *C5AR1* and *TLR2)* is most significantly enriched in WikiPathways gene sets “TYROBP Causal Network” and “Human Complement System” (**Figure 4E, Table S37**). GBA dysfunction has been associated with glucocerebroside-specific IgG autoantibodies, complement effector C5a generation and C5aR1 activation in a cycle that can drive innate and adaptive immune cell recruitment and activation in GD.^68,69^ We observed significant overlap between GD-associated modules and subnetworks and DEGs from recombinant C5a-treated human GD-induced pluripotent stem cell-derived macrophages (**Table S37,** “GD with C5a vs GD with vehicle”, “C5a all vs vehicle all”).^70^ Thus, we provide independent evidence for the association between GBA function, complement activation with *CTSS* upregulation, and GD-relevant phenotypes including increased liver mass and altered glycosphingolipids both in a model of GD and models of ‘normal’ variation and complex diseases. Recapitulation of these associations supports the biology and imbues confidence in the network approach to identify modifying pathways in IEM models.

## Discussion

In this study, we applied a novel approach that expands the search space for modifiers of IEM in a data-driven and unbiased way (**Figure 5A**). Several novel insights were gained. First, we added IEM-associated metabolites as BXD RI strain phenotypes for further investigation and identified a novel structural variant in the *Mlycd* DBA/2J haplotype causing a biochemical phenotype that may act as a modifier of IEM biology in the mouse. Second, we found that IEM disease signatures generated as DEGs from IEM mouse models (FAO disorders and GD) are coexpressed and highly connected in networks built from common population transcriptomes (human and mouse). This indicates that despite the extreme perturbation in the IEM condition, conserved biology is highlighted by smaller perturbations in the common population. We exploited this observation and identified and then experimentally validated GR signaling as a candidate modifier pathway that may explain differences in clinical presentation of FAO disorder patients. Using the same approach, we independently validated a link between complement signaling and inflammation in GD while also identifying a novel player in CTSS that may serve a role in translating the accumulation of glucosylceramide to a hyperinflammatory state. Our observations support finding modifiers of IEM through approaches that consider them as part of a continuum of disease states with the common population at the mild end of the extremes.^1^

**Figure 5.**
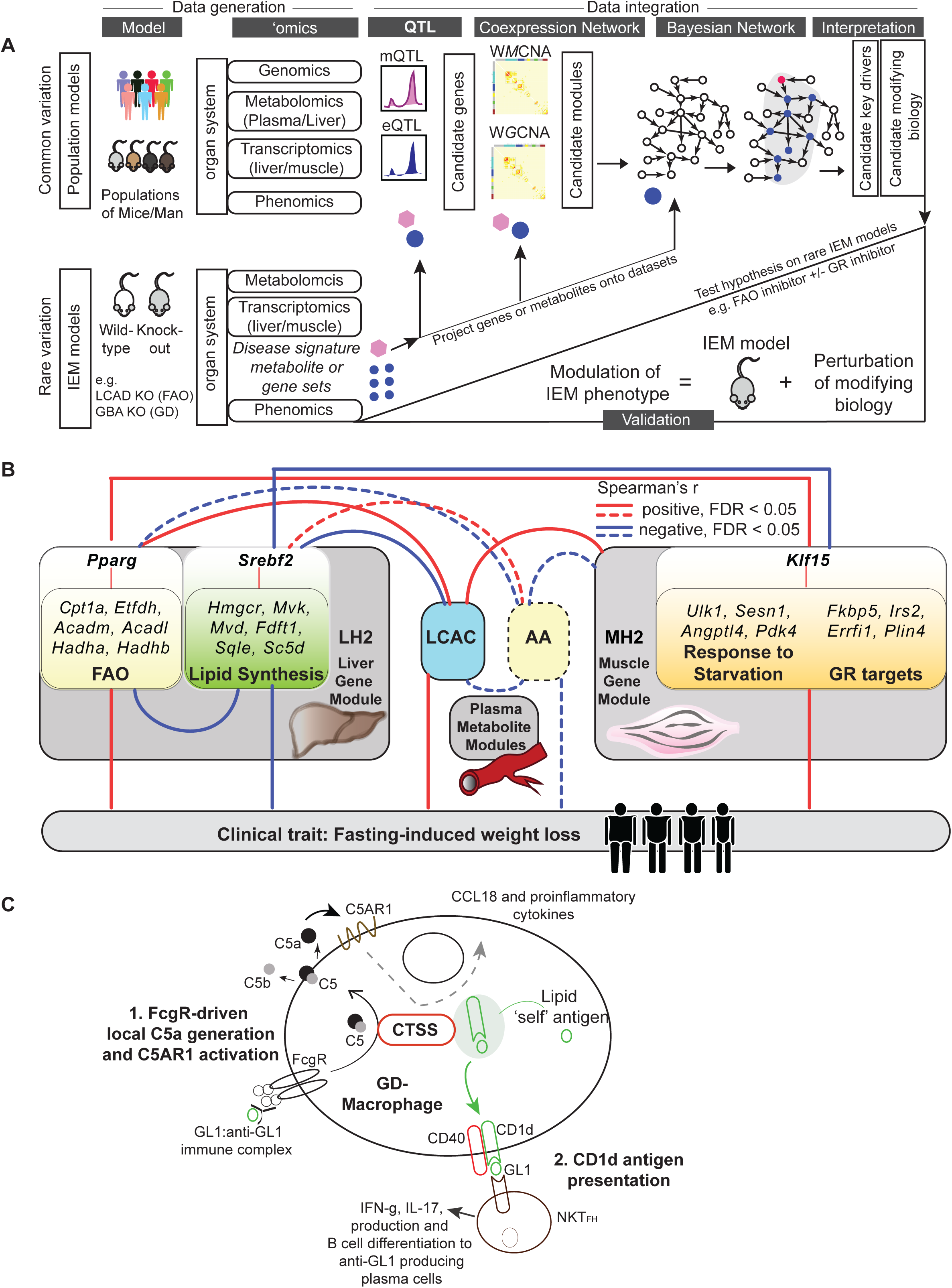
The multiomic network approach reveals a multi-tissue interaction of genes and metabolites that may underlie the phenotypic variability in IEM. (A) The multiomic network approach used to uncover disease modifying mechanisms of IEM. Schema that represents the approaches used to overcome the rare disease-rare data dilemma and find novel modifying biology of IEM. (B) Schema outlining the interactions between liver metabolism (FAO and cholesterol synthesis), muscle metabolism (role of GR activation in the starvation response), plasma metabolites (LCAC and AA) and clinical traits (fasting-induced weight loss). Population level variation in these interactions is speculated to act as a modifier of the pathophysiology of FAO disorders. (C) Schema outlining the potential impact of CTSS on complement activation and lipid self-antigen presentation by the MHC I-like CD1 family. Population level variation in these processes is speculated to act as modifiers of the pathophysiology of GD.

The LCAD KO mouse, a model for long-chain FAO disorders, displays increased LCAC, fatty liver, fasting-induced hypoketotic hypoglycemia and cardiac hypertrophy,^46–49^ which are similar phenotypes as observed in patients. Our prior work has demonstrated that impaired amino acid metabolism contributes to fasting-induced hypoglycemia in the LCAD KO mouse^48^ and impacts on metabolic signaling.^46^ In the BXD RI multiomic data, the liver and muscle gene expression modules we identified as most enriched for tissue-specific LCAD KO mouse DEGs correlated with LCAC, total AA and FIWL. It is known that during fasting, FAO serves as an alternative to glucose for ATP production. Being able to spare glucose is crucial to protect the body against excessive erosion of protein stores, since AAs are the main alternative source of gluconeogenic precursors.^71–73^ Indeed, it was recently demonstrated that liver alanine conversion into glucose can promote skeletal muscle atrophy.^74^ The current work further establishes the tight link between AA metabolism and mitochondrial FAO. Our data show for the first time that there is a genetic basis for the molecular coordination of the fasting response between liver and muscle, which may underlie some of the phenotypic variation observed in patients with a FAO disorder (**Figure 5B**, **Table S42**).

We further investigated the muscle LCAD KO DEG signature and its enriched coexpression modules in the context of muscle GRNs. Key driver analysis identified GR and KLF15 signaling as a pathway that may modulate the activity of these subnetworks and thus potentially underlie the variation in FIWL, LCAC and total AA. We inhibited GR signaling in a model for mitochondrial FAO deficiency and demonstrated reduced FIWL and more pronounced hypoglycemia. This result indicates that muscle GR signaling may be a candidate modifier of the phenotypes in long-chain FAO disorders. At this point, we are not able to pinpoint a genetic cause underlying variation in muscle GR signaling. This could be a muscle factor such as the key driver *Klf15*, but it is equally well conceivable that the cause is polygenic involving multiple tissue types. Glucocorticoid signaling is controlled by the hypothalamic–pituitary–adrenal axis, a neuroendocrine system that is influenced by stress, but also hypoglycemia. In accordance, the increase in glucose production is a classic function of glucocorticoids.

To illustrate that our method of integrating multi-scale data from general populations to identify modifying mechanisms in rare disease can be applied to other IEM, we studied GD-specific subnetworks. Our pathway enrichment and key driver analysis identified components of complement signaling as important in GD-associated pathology. As the expression of these genes was also correlated with the liver mass phenotype of the BXD RI cohort, it suggests complement-associated genes may underlie some of the associated liver pathology in GD. Indeed, activation of C5a and C5a receptor 1 (C5aR1) controls glucocerebroside accumulation and the inflammatory response in experimental and clinical GD.^68^ Furthermore, the GD mouse liver DEGs, GD-associated BXD modules LC4, LH3, and LH4, and GD-associated subnetworks are all enriched in gene sets generated from the treatment of C5a-treated, GD-patient derived macrophages,^70^ which further supports the link between these GD-specific subnetworks and complement activation. This recapitulation of complement as a driver in GD illustrates that our approach is generalizable and can be applied to other IEM.

*CTSS* was identified as a key driver of this GD pathology. CTSS is a member of the cysteine family of lysosomal proteases, predominately expressed in antigen-presenting cells and upregulated in models of antigen-induced inflammation.^75–79^ CTSS participates in the degradation of antigenic proteins to peptides for presentation on MHC class II and the degradation of antigenic lipids for presentation on CD1d.^80^ Further studies are required to determine if CTSS cleaves accumulated glucosylceramide for cell surface presentation by CD1d. In this case, recognition of CD1d-presented glucosylceramide fragments by NKT cells would drive B cell differentiation to anti-glucosylceramide autoantibody-producing plasma cells and a hyper-inflammatory state (**Figure 5C**). This hypothesis is supported by study of a *Ctss*^−/−^ model with a mild immunocompromised phenotype.^79^

Although CTSS itself has not been associated with complement signaling yet, other members of the cathepsin protease family have previously been implicated in this process. The related cathepsin H has been shown to cleave intracellular C5 to generate the potent chemotaxin C5a in extracellular compartments of atherosclerotic plaques.^81^ In addition, the related cathepsin L can process complement C3 into biologically active C3a and C3b in T cells.^82^ Importantly, intracellular generation of C3a was observed in not just T cells, but other immune and non-immune cell populations as well, suggesting that intracellular complement activation might be of broad physiological significance. While binding of the glucosylceramide-IgG auto-immune complexes to FcgR on phagocytes and antigen-presenting cells is known to lead to local C5a generation through C5 proteolysis, more study is required to determine if CTSS can act as this FcgR-activated protease. If this were true, then modulation of C5aR1 and its downstream effectors including CCL18 and other proinflammatory cytokines by CTSS would represent a novel complement-activating mechanism (**Figure 5C**). Even in the absence of the specific mechanism, we not only provide further evidence that connects complement signaling and the hyperinflammatory state associated with macrophage dysfunction and lipid dysregulation characteristic of GD, but we also identified a link between CTSS and complement signaling that may underlie diverse GD severity.

In summary, we demonstrate that the gene-gene interactions found within GRNs built from common disease transcriptomes can be exploited to derive IEM associated subnetworks. This provides new gene sets and key driver genes to be further annotated, characterized, and probed for modifying biology that connects primary disease genes with the clinical phenotype. We speculate that from these subnetworks, variants in new candidate modifiers will emerge as novel players in disease pathology as well as therapeutic targets to complement existing interventions.

## Supporting information

Supplementary Materials and Methods, Supplementary Figures and legends, Overview of all Supplementary Tables

## List of abbreviations

B6: C57BL/6J
BXD: cross of C57BL/6J and DBA/2J
C3DC: malonylcarnitine
CD: chow diet
D2: DBA/2J
DEG: differentially expressed genes
FAO: fatty acid oxidation
FIWL: fastinginduced weight loss
GBA: β-glucocerebrosidase
GRN: gene regulatory network
HFD: high fat diet
IEM: inborn errors of metabolism
LCAC: long-chain acylcarnitines
LCAD: long-chain acyl-CoA dehydrogenase
MCD: malonyl-CoA decarboxylase
RI: recombinant inbred
QTL: quantitative trait locus
SNP: single nucleotide polymorphism

## Declarations

## Acknowledgements

We acknowledge the help of the shared resource facilities at the Icahn School of Medicine at Mount Sinai (Colony Management, Real Time Polymerase Chain Reaction (qPCR), Mouse Genetics and Gene Targeting, Scientific Computing and the Genomics Core). DNA from SM/J mice was a kind gift from Dr. Weibin Shi (University of Virginia). The authors thank Simone Denis (Laboratory Genetic Metabolic Diseases) for tissue acylcarnitine analysis.

## Ethics approval

The animal research described in this study was approved by the Institutional Animal Care and Use Committee (IACUC) of the Icahn School of Medicine at Mount Sinai or by the Swiss cantonal veterinary authorities of Vaud under licenses 2257.0 and 2257.1.

## Authors’ contributions

SMH and CAA conceived this study. AB, SMH and CAA analyzed and interpreted all the data and were major contributors in writing the manuscript. PRR analyzed and interpreted the mifepristone/L-aminocarnitine data. EGW, MM, JAH, CNL, RJW, JMA, JZ and JA contributed essential methods, tools and/or models for this work. All authors read and approved the final manuscript.

## References

1 Argmann CA, Houten SM, Zhu J & Schadt EE. A Next Generation Multiscale View of Inborn Errors of Metabolism. Cell Metab 23, 13–26, doi:10.1016/j.cmet.2015.11.012 (2016).

2 Dipple KM & McCabe ER. Phenotypes of patients with “simple” Mendelian disorders are complex traits: thresholds, modifiers, and systems dynamics. Am J Hum Genet 66, 1729–1735, doi:10.1086/302938 (2000).

3 Scriver CR & Waters PJ. Monogenic traits are not simple: lessons from phenylketonuria. Trends in genetics : TIG 15, 267–272 (1999).

4 Dipple KM & McCabe ER. Modifier genes convert “simple” Mendelian disorders to complex traits. Mol Genet Metab 71, 43–50, doi:10.1006/mgme.2000.3052 (2000).

5 Surendran P, Stewart ID, Au Yeung VPW et al. Rare and common genetic determinants of metabolic individuality and their effects on human health. Nat Med, doi:10.1038/s41591-022-02046-0 (2022).

6 Shin SY, Fauman EB, Petersen AK et al. An atlas of genetic influences on human blood metabolites. Nat Genet 46, 543–550, doi:10.1038/ng.2982 (2014).

7 Scherer N, Fassler D, Borisov O et al. Coupling metabolomics and exome sequencing reveals graded effects of rare damaging heterozygous variants on gene function and human traits. Nat Genet, doi:10.1038/s41588-024-01965-7 (2025).

8 Belbin GM, Rutledge S, Dodatko T et al. Leveraging health systems data to characterize a large effect variant conferring risk for liver disease in Puerto Ricans. Am J Hum Genet 108, 2099–2111, doi:10.1016/j.ajhg.2021.09.016 (2021).

9 Barton AR, Hujoel MLA, Mukamel RE, Sherman MA & Loh PR. A spectrum of recessiveness among Mendelian disease variants in UK Biobank. Am J Hum Genet 109, 1298–1307, doi:10.1016/j.ajhg.2022.05.008 (2022).

10 Lupski JR, Belmont JW, Boerwinkle E & Gibbs RA. Clan genomics and the complex architecture of human disease. Cell 147, 32–43, doi:10.1016/j.cell.2011.09.008 (2011).

11 Blair DR, Lyttle CS, Mortensen JM et al. A nondegenerate code of deleterious variants in Mendelian loci contributes to complex disease risk. Cell 155, 70–80, doi:10.1016/j.cell.2013.08.030 (2013).

12 Schadt EE. Molecular networks as sensors and drivers of common human diseases. Nature 461, 218–223, doi:nature08454 [pii];10.1038/nature08454 [doi] (2009).

13 Di Narzo AF, Houten SM, Kosoy R et al. Integrative analysis of the inflammatory bowel disease serum metabolome improves our understanding of genetic etiology and points to novel putative therapeutic targets. Gastroenterology 162, 828–843, doi:10.1053/j.gastro.2021.11.015 (2022).

14 Argmann C, Tokuyama M, Ungaro RC et al. Molecular Characterization of Limited Ulcerative Colitis Reveals Novel Biology and Predictors of Disease Extension. Gastroenterology 161, 1953–1968 e1915, doi:10.1053/j.gastro.2021.08.053 (2021).

15 Chen Y, Zhu J, Lum PY et al. Variations in DNA elucidate molecular networks that cause disease. Nature 452, 429–435, doi:nature06757 [pii];10.1038/nature06757 [doi] (2008).

16 Peters LA, Perrigoue J, Mortha A et al. A functional genomics predictive network model identifies regulators of inflammatory bowel disease. Nat Genet 49, 1437–1449, doi:10.1038/ng.3947 (2017).

17 Zhu J, Sova P, Xu Q et al. Stitching together multiple data dimensions reveals interacting metabolomic and transcriptomic networks that modulate cell regulation. PLoS biology 10, e1001301, doi:10.1371/journal.pbio.1001301 (2012).

18 Yoo S, Takikawa S, Geraghty P et al. Integrative analysis of DNA methylation and gene expression data identifies EPAS1 as a key regulator of COPD. PLoS Genet 11, e1004898, doi:10.1371/journal.pgen.1004898 (2015).

19 Tu Z, Argmann C, Wong KK et al. Integrating siRNA and protein-protein interaction data to identify an expanded insulin signaling network. Genome Res. 19, 1057–1067 (2009).

20 Davis RC, van Nas A, Castellani LW et al. Systems genetics of susceptibility to obesity-induced diabetes in mice. Physiol Genomics 44, 1–13, doi:10.1152/physiolgenomics.00003.2011 (2012).

21 Schadt EE, Lamb J, Yang X et al. An integrative genomics approach to infer causal associations between gene expression and disease. Nat. Genet 37, 710–717, doi:ng1589 [pii];10.1038/ng1589 [doi] (2005).

22 Li Q, Lee CH, Peters LA et al. Variants in TRIM22 That Affect NOD2 Signaling Are Associated With Very-Early-Onset Inflammatory Bowel Disease. Gastroenterology 150, 1196–1207, doi:10.1053/j.gastro.2016.01.031 (2016).

23 Wang H, Bender A, Wang P et al. Insights into beta cell regeneration for diabetes via integration of molecular landscapes in human insulinomas. Nature communications 8, 767, doi:10.1038/s41467-017-00992-9 (2017).

24 Zhang CK, Stein PB, Liu J et al. Genome-wide association study of N370S homozygous Gaucher disease reveals the candidacy of CLN8 gene as a genetic modifier contributing to extreme phenotypic variation. American journal of hematology 87, 377–383, doi:10.1002/ajh.23118 (2012).

25 Ashbrook DG, Arends D, Prins P et al. A platform for experimental precision medicine: The extended BXD mouse family. Cell Syst 12, 235–247 e239, doi:10.1016/j.cels.2020.12.002 (2021).

26 Peirce JL, Lu L, Gu J, Silver LM & Williams RW. A new set of BXD recombinant inbred lines from advanced intercross populations in mice. BMC genetics 5, 7, doi:10.1186/1471-2156-5-7 (2004).

27 Williams EG, Wu Y, Jha P et al. Systems proteomics of liver mitochondria function. Science 352, aad0189, doi:10.1126/science.aad0189 (2016).

28 Wu Y, Williams EG, Dubuis S et al. Multilayered genetic and omics dissection of mitochondrial activity in a mouse reference population. Cell 158, 1415–1430, doi:10.1016/j.cell.2014.07.039 (2014).

29 Andreux PA, Williams EG, Koutnikova H et al. Systems genetics of metabolism: the use of the BXD murine reference panel for multiscalar integration of traits. Cell 150, 1287–1299, doi:S0092-8674(12)01007-0 [pii];10.1016/j.cell.2012.08.012 [doi] (2012).

30 Broman KW, Wu H, Sen S & Churchill GA. R/qtl: QTL mapping in experimental crosses. Bioinformatics 19, 889–890, doi:10.1093/bioinformatics/btg112 (2003).

31 Zhang B & Horvath S. A general framework for weighted gene co-expression network analysis. Statistical applications in genetics and molecular biology 4, Article17, doi:10.2202/1544-6115.1128 (2005).

32 Schadt EE, Molony C, Chudin E et al. Mapping the genetic architecture of gene expression in human liver. PLoS.Biol. 6, e107, doi:07-PLBI-RA-4030 [pii];10.1371/journal.pbio.0060107 [doi] (2008).

33 Zhu J, Lum PY, Lamb J et al. An integrative genomics approach to the reconstruction of gene networks in segregating populations. Cytogenetic and genome research 105, 363–374, doi:10.1159/000078209 (2004).

34 Zhang B & Zhu J. in The World Congress on Engineering. (eds S.I. Ao et al.) 1309–1312 (Newswood Limited).

35 Janky R, Verfaillie A, Imrichova H et al. iRegulon: from a gene list to a gene regulatory network using large motif and track collections. PLoS computational biology 10, e1003731, doi:10.1371/journal.pcbi.1003731 (2014).

36 Leandro J, Violante S, Argmann CA et al. Mild inborn errors of metabolism in commonly used inbred mouse strains. Mol Genet Metab 126, 388–396, doi:10.1016/j.ymgme.2019.01.021 (2019).

37 Ghazalpour A, Bennett BJ, Shih D et al. Genetic regulation of mouse liver metabolite levels. Molecular systems biology 10, 730, doi:10.15252/msb.20135004 (2014).

38 Salomons GS, Jakobs C, Pope LL et al. Clinical, enzymatic and molecular characterization of nine new patients with malonyl-coenzyme A decarboxylase deficiency. J Inherit Metab Dis 30, 23–28, doi:10.1007/s10545-006-0514-6 (2007).

39 Sun XY, Chen ZY, Hayashi Y et al. Insertion of an intracisternal A particle retrotransposon element in plasma membrane calcium ATPase 2 gene attenuates its expression and produces an ataxic phenotype in joggle mutant mice. Gene 411, 94–102, doi:10.1016/j.gene.2008.01.013 (2008).

40 Mietz JA, Grossman Z, Lueders KK & Kuff EL. Nucleotide sequence of a complete mouse intracisternal A-particle genome: relationship to known aspects of particle assembly and function. J Virol 61, 3020–3029 (1987).

41 Fabisiak JP, Medvedovic M, Alexander DC et al. Integrative metabolome and transcriptome profiling reveals discordant energetic stress between mouse strains with differential sensitivity to acrolein-induced acute lung injury. Molecular nutrition & food research 55, 1423–1434, doi:10.1002/mnfr.201100291 (2011).

42 Petkov PM, Graber JH, Churchill GA et al. Evidence of a large-scale functional organization of mammalian chromosomes. PLoS Genet 1, e33, doi:10.1371/journal.pgen.0010033 (2005).

43 Soeters MR, Serlie MJ, Sauerwein HP et al. Characterization of D-3-hydroxybutyrylcarnitine (ketocarnitine): an identified ketosis-induced metabolite. Metabolism 61, 966–973, doi:10.1016/j.metabol.2011.11.009 (2012).

44 McGarry JD, Mannaerts GP & Foster DW. A possible role for malonyl-CoA in the regulation of hepatic fatty acid oxidation and ketogenesis. J. Clin. Invest. 60, 265–270, doi:10.1172/JCI108764 (1977).

45 Drynan L, Quant PA & Zammit VA. Flux control exerted by mitochondrial outer membrane carnitine palmitoyltransferase over beta-oxidation, ketogenesis and tricarboxylic acid cycle activity in hepatocytes isolated from rats in different metabolic states. Biochem. J. 317 (Pt 3), 791–795 (1996).

46 Ranea-Robles P, Pavlova NN, Bender A et al. A mitochondrial long-chain fatty acid oxidation defect leads to tRNA uncharging and activation of the integrated stress response in the mouse heart. Cardiovascular research, doi:10.1093/cvr/cvac050 (2022).

47 Kurtz DM, Rinaldo P, Rhead WJ et al. Targeted disruption of mouse long-chain acyl-CoA dehydrogenase gene reveals crucial roles for fatty acid oxidation. Proc. Natl. Acad. Sci. USA 95, 15592–15597 (1998).

48 Houten SM, Herrema H, te Brinke H et al. Impaired amino acid metabolism contributes to fasting-induced hypoglycemia in fatty acid oxidation defects. Hum. Mol. Genet. 22, 5249–5261, doi:10.1093/hmg/ddt382 (2013).

49 Bakermans AJ, Geraedts TR, van Weeghel M et al. Fasting-induced myocardial lipid accumulation in long-chain acyl-CoA dehydrogenase knock-out mice is accompanied by impaired left ventricular function. Circ. Cardiovasc. Imaging 4, 558–565, doi:CIRCIMAGING.111.963751 [pii];10.1161/CIRCIMAGING.111.963751 [doi] (2011).

50 Diekman EF, van Weeghel M, Wanders RJ, Visser G & Houten SM. Food withdrawal lowers energy expenditure and induces inactivity in long-chain fatty acid oxidation-deficient mouse models. FASEB J. 28, 2891–2900, doi:10.1096/fj.14-250241 (2014).

51 GTEx Consortium. Human genomics. The Genotype-Tissue Expression (GTEx) pilot analysis: multitissue gene regulation in humans. Science 348, 648–660, doi:10.1126/science.1262110 (2015).

52 Kuo T, Lew MJ, Mayba O et al. Genome-wide analysis of glucocorticoid receptor-binding sites in myotubes identifies gene networks modulating insulin signaling. Proc Natl Acad Sci USA 109, 11160–11165, doi:10.1073/pnas.1111334109 (2012).

53 Fan L, Sweet DR, Prosdocimo DA et al. Muscle Kruppel-like factor 15 regulates lipid flux and systemic metabolic homeostasis. J Clin Invest 131, e139496, doi:10.1172/JCI139496 (2021).

54 Sasse SK, Mailloux CM, Barczak AJ et al. The glucocorticoid receptor and KLF15 regulate gene expression dynamics and integrate signals through feed-forward circuitry. Mol Cell Biol 33, 2104–2115, doi:10.1128/MCB.01474-12 (2013).

55 Jeyaraj D, Scheer FA, Ripperger JA et al. Klf15 orchestrates circadian nitrogen homeostasis. Cell Metab 15, 311–323, doi:10.1016/j.cmet.2012.01.020 (2012).

56 Gray S, Wang B, Orihuela Y et al. Regulation of gluconeogenesis by Kruppel-like factor 15. Cell Metab 5, 305–312, doi:10.1016/j.cmet.2007.03.002 (2007).

57 Ranea-Robles P, Violante S, Argmann C et al. Murine deficiency of peroxisomal L-bifunctional protein (EHHADH) causes medium-chain 3-hydroxydicarboxylic aciduria and perturbs hepatic cholesterol homeostasis. Cellular and molecular life sciences : CMLS 78, 5631–5646, doi:10.1007/s00018-021-03869-9 (2021).

58 Violante S, Achetib N, van Roermund CWT et al. Peroxisomes can oxidize medium- and long-chain fatty acids through a pathway involving ABCD3 and HSD17B4. FASEB J 33, 4355–4364, doi:10.1096/fj.201801498R (2019).

59 Jenkins DL & Griffith OW. Antiketogenic and hypoglycemic effects of aminocarnitine and acylaminocarnitines. Proc Natl Acad Sci USA 83, 290–294 (1986).

60 Chegary M, te Brinke H, Doolaard M, et al. Characterization of L-aminocarnitine, an inhibitor of fatty acid oxidation. Mol. Genet. Metab. 93, 403–410 (2008).

61 Dasgupta N, Xu YH, Oh S et al. Gaucher disease: transcriptome analyses using microarray or mRNA sequencing in a Gba1 mutant mouse model treated with velaglucerase alfa or imiglucerase. PloS one 8, e74912, doi:10.1371/journal.pone.0074912 (2013).

62 Schadt EE, Molony C, Chudin E et al. Mapping the genetic architecture of gene expression in human liver. PLoS biology 6, e107, doi:10.1371/journal.pbio.0060107 (2008).

63 Yang X, Zhang B, Molony C et al. Systematic genetic and genomic analysis of cytochrome P450 enzyme activities in human liver. Genome Res. 20, 1020–1036, doi:gr.103341.109 [pii];10.1101/gr.103341.109 [doi] (2010).

64 Jmoudiak M & Futerman AH. Gaucher disease: pathological mechanisms and modern management. Br J Haematol 129, 178–188, doi:10.1111/j.1365-2141.2004.05351.x (2005).

65 Mistry PK, Taddei T, vom Dahl S & Rosenbloom BE. Gaucher disease and malignancy: a model for cancer pathogenesis in an inborn error of metabolism. Crit Rev Oncog 18, 235–246, doi:10.1615/critrevoncog.2013006145 (2013).

66 Sidransky E, Nalls MA, Aasly JO et al. Multicenter analysis of glucocerebrosidase mutations in Parkinson’s disease. N Engl J Med 361, 1651–1661, doi:10.1056/NEJMoa0901281 (2009).

67 Afinogenova Y, Ruan J, Yang R et al. Aberrant progranulin, YKL-40, cathepsin D and cathepsin S in Gaucher disease. Mol Genet Metab 128, 62–67, doi:10.1016/j.ymgme.2019.07.014 (2019).

68 Pandey MK, Burrow TA, Rani R et al. Complement drives glucosylceramide accumulation and tissue inflammation in Gaucher disease. Nature 543, 108–112, doi:10.1038/nature21368 (2017).

69 Pandey MK, Grabowski GA & Kohl J. An unexpected player in Gaucher disease: The multiple roles of complement in disease development. Semin Immunol 37, 30–42, doi:10.1016/j.smim.2018.02.006 (2018).

70 Serfecz JC, Saadin A, Santiago CP et al. C5a Activates a Pro-Inflammatory Gene Expression Profile in Human Gaucher iPSC-Derived Macrophages. Int J Mol Sci 22, doi:10.3390/ijms22189912 (2021).

71 Owen OE, Smalley KJ, D’Alessio DA, Mozzoli MA & Dawson EK. Protein, fat, and carbohydrate requirements during starvation: anaplerosis and cataplerosis. Am. J. Clin. Nutr. 68, 12–34 (1998).

72 Nurjhan N, Bucci A, Perriello G et al. Glutamine: a major gluconeogenic precursor and vehicle for interorgan carbon transport in man. J. Clin. Invest. 95, 272–277, doi:10.1172/JCI117651 [doi] (1995).

73 Cahill GF, Jr. Fuel metabolism in starvation. Annu. Rev. Nutr. 26, 1–22, doi:10.1146/annurev.nutr.26.061505.111258 (2006).

74 Okun JG, Rusu PM, Chan AY et al. Liver alanine catabolism promotes skeletal muscle atrophy and hyperglycaemia in type 2 diabetes. Nat Metab 3, 394–409, doi:10.1038/s42255-021-00369-9 (2021).

75 Beers C, Burich A, Kleijmeer MJ et al. Cathepsin S controls MHC class II-mediated antigen presentation by epithelial cells in vivo. J Immunol 174, 1205–1212, doi:10.4049/jimmunol.174.3.1205 (2005).

76 Dheilly E, Battistello E, Katanayeva N et al. Cathepsin S Regulates Antigen Processing and T Cell Activity in Non-Hodgkin Lymphoma. Cancer Cell 37, 674–689 e612, doi:10.1016/j.ccell.2020.03.016 (2020).

77 Costantino CM, Ploegh HL & Hafler DA. Cathepsin S regulates class II MHC processing in human CD4+ HLA-DR+ T cells. J Immunol 183, 945–952, doi:10.4049/jimmunol.0900921 (2009).

78 Riese RJ, Mitchell RN, Villadangos JA et al. Cathepsin S activity regulates antigen presentation and immunity. J Clin Invest 101, 2351–2363, doi:10.1172/JCI1158 (1998).

79 Thanei S, Theron M, Silva AP et al. Cathepsin S inhibition suppresses autoimmune-triggered inflammatory responses in macrophages. Biochem Pharmacol 146, 151–164, doi:10.1016/j.bcp.2017.10.001 (2017).

80 de Mingo Pulido A, de Gregorio E, Chandra S et al. Differential Role of Cathepsins S and B In Hepatic APC-Mediated NKT Cell Activation and Cytokine Secretion. Front Immunol 9, 391, doi:10.3389/fimmu.2018.00391 (2018).

81 Perez HD, Ohtani O, Banda D et al. Generation of biologically active, complement-(C5) derived peptides by cathepsin H. J Immunol 131, 397–402 (1983).

82 Liszewski MK, Kolev M, Le Friec G et al. Intracellular complement activation sustains T cell homeostasis and mediates effector differentiation. Immunity 39, 1143–1157, doi:10.1016/j.immuni.2013.10.018 (2013).

