## Supplementary Materials and Methods, Supplementary Figures and legends, Overview of all Supplementary Tables for "A multiomic network approach uncovers disease modifying mechanisms of inborn errors of metabolism"

*BXD RI strains*

We phenotyped 42 RI strains from the BXD population [1, 2]. Phenotypes and genotypes are available from www.genenetwork.org. From each strain, ~10 male animals were selected and separated at 8 weeks of age into two equal groups for each dietary cohort. Mice were fed a chow diet (CD, Harlan Teklad Global 18% Protein Rodent Diet, 2018, 6% kcal/fat) or a high fat diet (HFD, Harlan Teklad, TD.06414, 60% kcal/fat). One strain did not give sufficient male pups for both cohorts, leaving 42 CD strains and 41 HFD strains, of which 41 overlap. The diet challenge was implemented until sacrifice at 29 weeks of age. All animals within each cohort were co-housed until week 23 of age, after which animals were single housed. At week 29, animals were fasted overnight, anesthetised using isoflurane and euthanized by exsanguination.

Fasting-induced weight loss (FIWL) refers to trait IDs 17609 (CD) and 17610 (HFD) and is the weight in g an animal loses during the overnight fast directly before the necropsy.

*Sample preparation*

Sample generation was described previously [1, 2]. Briefly, for plasma isolation, approximately 1 mL was collected from the vena cava, 800 µL of which was placed in lithium-heparin coated tubes then centrifuged at 4,000 rpm for 5 minutes at 4°C. The isolated plasma was flash frozen in liquid nitrogen and stored at -80°C for later analysis, which was done in tandem for all samples. For all cohorts, the plasma was pooled equally by volume for all animals to create sufficient quantities for all metabolite analyses. Sufficient plasma for metabolomics was collected from 42 RI strains of which 38 were samples in both diets. Fourteen strains had sufficient plasma to analyze both individuals and pooled cohorts (8 CD, 6 HFD), allowing for the calculation of the heritability estimates by comparing within-strain variance versus across-strain and across-diet variance. For tissue collection, animals were perfused with PBS immediately following blood draw. The liver (gallbladder removed) and quadriceps (from both legs) were then dissected and flash frozen in liquid nitrogen.

*RNA isolation and microarray analysis*

For transcriptomic analysis, a ~100 mg piece of both liver and quadriceps tissue was used. mRNA for ~5 animals per cohort was isolated, and then pooled evenly (~8 µg of RNA for each mouse) into a single RNA sample for each cohort. These pooled RNA samples were then purified using

RNeasy (Qiagen) and analyzed using the Affymetrix Mouse Gene 1.0 ST array. All RIN values were > 8.0 [1, 2].

*Metabolite measurements*

Sixty-five plasma metabolites and 12 liver metabolites were measured in the BXD RI cohort. Plasma amino acids and acylcarnitines were measured using tandem mass spectrometry (MS/MS)[3-6]. Additional metabolites (glucose, HDL, LDL, and cholesterol) were curated from previous reports because of their dysregulation in different IEM and to help organize coexpressed metabolites [1, 2]. Plasma ceramide, sphingomyelins and glycosphingolipids were measured based on derivatization of the amine in the glycolipid with orthophthaldialdehyde (OPA), followed by high performance liquid chromatography (HPLC) separation and fluorescence detection [7]. Plasma phingosine-1-phosphate was measured by liquid chromatography MS/MS [8]. Globotriaosylsphingosine and gluco(galacto)sylsphingosine (lysoGb3) in liver were measured using LC-MS/MS [9]. All data are available on GeneNetwork.org under the “BXD Published Phenotypes” set.

*QTL assessment in the BXD RI cohorts*

Plasma and liver metabolite measurements were tested for normality by making QQ plots and doing the Shapiro-Wilk test (cut-off $p<0.001$). Measurements that were not normally distributed were natural log transformed (Phe, Arg, Pro, Asp, C3, C4, C5, C3DC, and N-glycolyl GM2). Metabolite measurements were then correlated with genotype for single nucleotide polymorphisms (SNPs) at 7,320 molecular markers in linkage disequilibrium across the genome using the Haley-Knott regression model of normally distributed traits using the software package R/qtl [10]. mQTL analysis was performed for metabolites measured in the CD cohort, the HFD cohort, and for combined traits that averaged the values of CD and HFD cohort measurements. These CD.HFD data points were aimed at increasing power of QTL detection by decreasing the variance in measurements. Transcriptome-wide eQTL analysis was performed on liver and muscle expression data for both dietary conditions separately (CD and HFD) using 25,199 probes per dataset. Microarray data were normalized using the “2Z+8” method in which gene-level Affymetrix 1.0 gene data were log transformed and centered around a value of 8 with a standard deviation of 2 [1, 2]. A non-parametric regression model was used to account for gene expression that is not normally distributed. Genome-scan adjusted p-values were calculated by permutation analysis (n = 1000), and QTLs with a log odds ratio (LOD) more extreme than at least 95% of permuted LOD values (corresponding to a genome-scan adjusted p-value < 0.05) were considered significant genome-wide. Gene probes with significant eQTLs within 1.5 MB of their start or end site were labeled *cis*-eQTLs, and broad *cis*-eQTL peaks were collapsed into lead SNPs according to lowest genome-scan adjusted p-value and highest LOD. Unique Affymetrix probes were mapped to Mouse Genome Informatics (MGI) gene symbols and human gene symbols from the HUGO Gene Nomenclature Committee (HGNC) sourced from the Ensembl database using the getLDS function of the biomaRt library in R (version 4.0.0).

*Prioritizing candidate causal genes in mQTL regions*

For metabolites with significant mQTLs, broad mQTL peaks were collapsed into single lead SNPs according to lowest genome-scan adjusted p-value (and highest LOD when necessary) per mQTL peak. Genes within mQTL regions were defined by Affymetrix probes starting or ending within 5 MB of the lead mQTL SNP. Genes within 5MB of lead mQTL SNPs were intersected with significant *cis*-eQTL data to define *cis*-eQTLs within mQTL regions. Metabolite levels were correlated to the expression of each gene with a significant eQTL (Spearman's rho) and the p value was adjusted for false discovery rate (FDR) using the Benjamini-Hochberg method.

*RNAseq of C57BL/6J and DBA/2J liver*

Samples from C57BL/6J and DBA/2J mice were collected previously to study naturally occurring biochemical traits in inbred mice [11, 12]. Total RNA from liver tissue of two C57BL/6J and two DBA/2J mice was isolated using QIAzol lysis reagent followed by purification using the RNeasy kit (Qiagen). RNA was submitted to the Genomics Core Facility at the Icahn School of Medicine at Mount Sinai for further processing. mRNA-focused cDNA libraries were generated using Illumina reagents (polyA capture), and samples were run on an Illumina HiSeq 2500 sequencer to yield a read depth of approximately 51 million 100 nucleotide single end reads per sample. Reads from fastq files were aligned to the mouse genome mm10 (GRCm38.75) with STAR release 2.4.0g1 [13] and summarized to gene- and exon-level counts using featureCounts version 1.5.2 [14]. The aligned reads and splice junctions were visualized using the Integrative Genomics Viewer (IGV) and its Sashimi plot feature [15, 16]. Sequencing data for C57BL/6J and DBA/2J liver samples were deposited in the GEO database (GSE186973).

*PacBio Single Molecule, Real-Time (SMRT) sequencing*

The retrotransposon in *Mlycd* was amplified using a forward primer in intron 2 (5'-gcc ctc tca tcc agt tgt aaa tgc tct ccg-3') and a reverse primer in exon 3 (5'-ggg tgg agc agt ggg aga aga agt aac atc-3'). Amplified material was prepared using the Amplicon Template Preparation and

Sequencing protocol and Pacific Biosciences Template Preparation Kit 1.0 according to manufacturer's instructions. SMRTbells were bound to P6 polymerase and sequenced using C4 chemistry and 360-minute movies on a single SMRTcell using the RSII system. Data were processed using the Long Amplicon Analysis pipeline contained within the SMRTPortal analysis suite (version 2.3.0) and resulted in a sequence of 6,043 bp in length and 99.994% accuracy (GenBank MH036232).

*Generation and characterization of a congenic DBA/2J line with Mlycd$^{B6/B6}$*

In the BXD6 strain, the C57BL/6J marker allele gnf08.119.598 was isolated in a DBA/2-context. Using speed congenics with the assistance of Jackson Laboratories, the BXD6 strain was crossed back in five generations to a DBA/2J background with exception of this single B6J-marker. The resulting line was named DBA/2J *Mlycd*$^{B6/B6}$ (D2.B6-*gnf08.119.598*).

The cohort for sample collection consisted of 9 DBA/2J mice (4 male and 5 female) and 6 DBA/2J *Mlycd*$^{B6/B6}$ (2 male and 4 female) with an average age of 24 weeks (range 17-27 weeks). Mice were subjected to overnight food withdrawal. Mice were anesthetized using pentobarbital (intraperitoneal 120 mg/kg in 0.9% NaCl) and then euthanized by exsanguination. The blood was collected from the vena cava inferior for the preparation of EDTA plasma. Organs were snap frozen in liquid nitrogen and stored at −80ºC. Liver acylcarnitine levels were measured as described [17, 18].

*RNAseq analysis of LCAD KO liver and gastrocnemius muscle*

Total RNA was isolated from liver and gastrocnemius muscle of 25 mice with the following genotypes (all male; 4 *Acadl*$^{+/+}$/*Acadvl*$^{+/+}$; 4 *Acadl*$^{+/-}$/*Acadvl*$^{+/+}$; 4 *Acadl*$^{+/-}$/*Acadvl*$^{+/-}$; 4 *Acadl*$^{-/-}$/*Acadvl*$^{+/+}$; 5 *Acadl*$^{-/-}$/*Acadvl*$^{+/-}$; 4 *Acadl*$^{+/-}$/*Acadvl*$^{-/-}$) described in Diekman et al [19]. RNA was submitted to the Genomics Core Facility at the Icahn School of Medicine at Mount Sinai for further processing. mRNA-focused cDNA libraries were generated using Illumina reagents (polyA capture), and samples were run on an Illumina HiSeq 2500 sequencer to yield a read depth of approximately 22 million 100 nucleotide single end mapped reads per muscle sample and approximately 33 million 100 nucleotide single end mapped reads per liver sample. Reads from fastq files were aligned to the mouse genome mm10 (GRCm38.75) with STAR release 2.3.0e [13] and summarized to gene level counts using featureCounts v1.4.4 [14]. Raw counts for all genes can be accessed on the Gene Expression Omnibus of the National Center for the Biotechnology Information for liver (GSE186613) and muscle (GSE186648) samples.

*Generation of rare disease signatures*

The LCAD KO signature was derived from a comparison of WT (*Acadl*$^{+/+}$/*Acadvl*$^{+/+}$) with LCAD KO mice (*Acadl*$^{-/-}$/*Acadvl*$^{+/+}$). Differential gene expression analysis (on 15,915 genes for liver and 13,394 genes for muscle) was conducted with the R package DESeq2 [20] as previously described [21]. Significant DEGs were defined using an adjusted p value < 0.05 (Benjamini-Hochberg method) with no fold change cut-off.

Publicly available, normalized liver gene expression data from a GD mouse model *Gba* p.D409V/null mice) was accessed and analyzed for DEGs using DESeq (FDR < 0.05, fold change cut-off of ±1.5). A molecular signature of GD was defined as the DEGs [22].

*Generation of coexpression networks from the BXD cohort*

Plasma metabolites with highly correlated expression values across samples were clustered into metabolite coexpression modules using an unsigned weighted gene coexpression analysis (WGCNA, beta = 7) [23]. Modules were named according to their main constituents; LCAC (turquoise), SCAC (blue), lipids (yellow), BCAA (green), and total AA (brown). To reduce data dimensionality from all constituent metabolite values to a single quantitative trait, the module eigenmetabolite was calculated as the first principal component of each module.

With the goal of identifying coexpressed genes across samples in the BXD cohort that also co-correlate with plasma and/or liver metabolites, liver and muscle genes were first independently organized into modules using WGCNA [23]. Unique Affymetrix probes, some mapping to the same gene, were collapsed to unique genes using the collapseRows function [24] and the connectivityBasedCollapsing argument of the WGCNA package in R. The module eigengene was calculated as the first principal component of each module.

BXD gene coexpression module eigengenes were correlated to plasma metabolite module eigenmetabolites, liver metabolite levels, and clinical trait expression levels assuming non-normal distribution of trait values (Spearman's r). P values were adjusted for multiple testing using the Benjamini-Hochberg method.

*Bayesian gene regulatory networks (GRNs)*

Mouse Bayesian GRNs were generated as previously described from the liver and muscle gene expression data generated from a series of segregating mouse populations [25]. These included the BXH/wt (C57BL/6J (B6) mice intercrossed with C3H/HeJ (C3H) mice to generate 321 F2 progeny (161 females, 160 males)) and the BXC crosses (C57BL/6J (B6) mice intercrossed with Castaneus (CAST) mice to generate 442 F2 progeny (276 females, 166 males)) [25]. The human liver GRN was constructed from the human liver cohort (HLC) comprised of 427 Caucasian subjects [25]. The muscle GRN was newly constructed as described [26] from muscle gene expression data obtained from the publicly available datasets on the Genotype-Tissue Expression (GTEx) portal, which contains data scored from 'normal' individuals across a spectrum of healthy to individuals with common diseases. The GTEx Project was supported by the Common Fund of the Office of the Director of the National Institutes of Health, and by NCI, NHGRI, NHLBI, NIDA, NIMH, and NINDS.

In short, transcript abundances and genotypes when available were input to the network reconstruction algorithm RIMBANET, and causal relationships were inferred between genes according to calculated posterior probabilities that variation in a given gene results in the perturbation of a second gene [26-28]. To simplify the exhaustive calculations of network construction, the specific constraint was implemented that a given gene could not be controlled by more than three 'parent' genes. The mouse liver network contains 9,338 nodes and 18,485 edges; the human liver network contains 8,336 nodes and 11,582 edges; the mouse muscle network contains 5,163 nodes and 8,622 edges and the human muscle network contains 8,458 nodes 16,422 edges.

*Generation of tissue-specific, disease-specific subnetworks*

Liver and muscle BXD coexpression modules were tested for enrichment in tissue-specific LCAD KO DEGs using a one-side Fisher's exact test. Similarly, liver modules were tested for enrichment in mutant *Gba* DEGs. P values were corrected for multiple testing using the Benjamini-Hochberg method. Individual tissue-specific, disease-specific subnetworks were generated by seeding LCAD KO or mutant *Gba* DEGs or their enriched BXD coexpression modules in tissue-matched GRNs. Unique Affymetrix identifiers from BXD array sequencing data, Ensembl gene IDs from LCAD KO sequencing data, and MGI gene symbols from curated mouse gene sets were mapped to orthologous human HGNC gene symbols via the useMart function of the biomaRt library in R [29]. Using the network structure, a variable number of layers (0-2) of nearest neighbor genes was included to generate the subnetworks (MCS). Subnetworks generated from the intersection of disease signatures or disease signature-enriched BXD gene coexpression modules were tested for enrichment in the original disease signatures using a one-sided Fisher's exact test with p values adjusted for multiple tests using the Benjamini and Hochberg method. Only subnetworks that remained enriched in the original seed signature upon incorporation of genes from the network structure were considered as most connected subnetworks (MCS) and used for downstream analysis.

*Shortest path analysis*

Disease signatures and signature-enriched BXD gene coexpression modules were evaluated for how well they were captured by tissue-specific GRNs. Using the distances function of the R package igraph, a matrix was generated to reflect the shortest path distance between every pair of nodes in the GRN. Genes that are directly connected have a shortest path distance of 1, genes separated by only one node between them have a shortest path distance of 2, and so on. The mean shortest path for all genes within each disease-associated gene set was calculated. For each gene set, 1,000 random gene sets of equal size were generated from the network, and the means of those random samples were used to generate a normalized distribution of random shortest path lengths. A normalized shortest path length (z-score) was then created for the disease-associated gene sets by comparing the mean shortest path lengths to their respective distributions. For example, 164 LCAD KO muscle DEGs were present in the human GRN, therefore the mean shortest path of 164 DEGs was compared to 1,000 mean shortest paths of random gene sets of size 164, and a z-score was generated. A normalized shortest path length less than -2 was considered to reflect a gene set that was more connected in the GRN than would be expected by random chance.

*Functional annotation of gene sets, coexpression modules and subnetworks*

BXD coexpression modules were tested for enrichment in genes from Gene Ontology (GO) biological pathways (version 2019-06-04) [30], Kyoto Encyclopedia of Genes and Genomes (KEGG) pathways (version available 2019-02-27) [31], and Reactome pathways (version available 2019-02-27) [32]. BXD coexpression modules enriched in LCAD KO DEGs and correlating to plasma LCAC levels, plasma AA levels, and fasting-induced weight loss were tested for enrichment in canonical gene sets with a two-sided hypergeometric test using the ClueGO plug-in version 2.5.4 for Cytoscape [33]. P values were corrected for multiple testing using the Bonferroni method. Most connected subnetworks associated with LCAD KO and GD were tested for enrichment in genes from the Molecular Signatures Database (MSigDB) hallmark gene set collection [34] reflecting chemical-gene interactions and disease signatures from the Comparative Toxicogenomic Database (CTD) [35] using a one-sided Fisher’s exact test. P values were corrected for multiple testing using the Benjamini-Hochberg method.

LCAD KO muscle DEGs, BXD coexpression modules enriched in LCAD KO muscle DEGs, and muscle-specific subnetworks generated from LCAD KO muscle DEGs and enriched modules were tested for enrichment in glucocorticoid receptor target genes and *Klf15* KO DEGs using a one-side Fisher's exact test. P values were corrected for multiple testing using the Benjamini-Hochberg method. The glucocorticoid receptor target gene set was curated from Kuo et al [36] and included 587 DEGs (367 upregulated, 220 downregulated; FDR < 0.05, fold change > 1.5) from cultured C2C12 myotubes treated with the GR agonist dexamethasone. Of these 587 DEGs, 173 genes (147 induced, 26 repressed) were determined to contain a glucocorticoid receptor binding region via ChIP-seq analysis. Annotation of key drivers of LCAD KO-associated subnetworks also included other literature supported GR targets and NR3C1 targets predicted by iRegulon [37]. A *Klf15* KO gene set was defined as the DEGsfrom quadriceps skeletal muscle tissue of a *Klf15*$^{-/-}$ mouse model following an 18 hour fast [38]. Gene expression data from GEO dataset GSE7137 were accessed using GEOquery (version 2.40.0) in R. Differential expression analysis was performed using limma (version 3.26.8). Genes with a nominal p-value < 0.01 were included in the *Klf15* KO skeletal muscle KO signature.

GD-related signatures, associated BXD coexpression modules, and associated subnetworks were tested for enrichment in gene sets generated from differential expression analysis of macrophages derived from GD patients treated with complement signaling agonist C5a (significant genes with FDR < 0.05) [39].

*Key driver analysis (KDA)*

Key driver analysis was performed using the KDA library in R [40]. Input gene sets were defined by subnetworks generated from disease-specific signatures (LCAD KO or GD) or subnetworks generated from signature-enriched BXD gene coexpression modules upon intersection with tissue-specific GRNs (mouse or human). For LCAD KO-associated liver KDA, generation of the LCAD KO DEG-associated subnetworks was further restricted by using only genes that were both differentially expressed with an FDR < 0.05 and whose mean expression level in LCAD KO samples was either doubled or halved compared to WT. Each subnetwork was independently tested as a gene set. For each node in the subnetwork, the enrichment was tested in its gene neighborhood within k steps (k varies from 1 to K) downstream and upstream for that individual gene set. Here, we defined K = 2 for all DEG- and module-based subnetworks.

*Transcription factor binding motif enrichment: iRegulon*

Disease-specific, tissue-specific subnetwork genes were tested for enrichment in canonical regulatory motifs involved in transcriptional regulation using ChIP-seq derived gene sets in iRegulon [37]. The enriched motifs were clustered by similarity, and motifs that belong to the same cluster were given the same cluster code. The cluster was assigned according to the maximal normalized enrichment score per motif cluster. Once enriched motifs were identified, an algorithm was applied to predict which transcription factors (TFs) are likely to bind those motifs. A maximum false discovery rate (FDR) for the prediction of TF binding to the motif was reported, with perfect matches labeled as 'Direct'. Predicted TFs were then annotated by iRegulon with known downstream genes.

*Validation of the role of glucocorticoid signaling during food withdrawal in a mouse model with a FAO defect*

The cohort for the L-AC (L-aminocarnitine [41, 42]) and/or mifepristone treatment consisted of 7 vehicle-treated (5 males and 2 females), 7 L-AC-treated (5 males and 2 females), 7 mifepristone-treated (5 males and 2 females), and 7 L-AC and mifepristone-treated (5 males and 2 females) 2-4 month-old mice. L-AC ((*R*)-aminocarnitine, minimum 97%) was obtained from Toronto Research Chemical Inc. (Toronto, ON, Canada). Mifepristone (RU-486, ≥98%) was obtained from Sigma (M8046). For mifepristone treatment, 2 mg of mifepristone were first dissolved in 0.04 mL of ethanol (50 mg/mL solution), followed by adding 0.025 mL of sterile 10% PEG-300/8% ethanol and 0.025 mL of sterile Tween-80. This solution was further diluted by adding 0.91 mL of sterile 0.9% NaCl. Using this cosolvent, mifepristone (2 mg/mL) remains soluble for several hours. Mice received an intraperitoneal injection of vehicle (cosolvent solution in 0.9% NaCl), or mifepristone (20 mg/kg) at 4:30 p.m. One hour later, mice received an intraperitoneal injection of vehicle (0.9% NaCl), or L-AC (16 mg/kg), followed by overnight food withdrawal. Blood glucose was measured after overnight food withdrawal using Bayer Contour blood glucose strips. Plasma corticosterone was measured using the Corticosterone Enzyme Immunoassay Kit (Arbor Assays, Cat #K014-H1). Mice were euthanized by exposure to $CO_2$ and blood was collected from the inferior vena cava for the preparation of EDTA plasma. Organs were snap frozen in liquid nitrogen and stored at −80ºC. *Klf15* mRNA was measured by qRT-PCR using primers (forward: GCGAGAAGCCC-TTTGCCT, reverse: GCTTCACACCCGAGTGAGAT).

**Supplementary Figure legends**

**Figure S1. Identifying potential modifiers of IEM metabolite abundance through QTL mapping.** (A) Unsupervised hierarchical clustering of the abundance of all metabolites measured in 40 BXD RI strains in two dietary conditions: chow diet (CD) and high fat diet (HFD). (B) The significant negative correlation between plasma C3DC levels and *Mlycd* gene expression in the liver of BXD mice on CD. The animals almost perfectly separated expression levels according to haplotype near the *Mlycd* gene. Animals with the DBA/2J haplotype had low *Mlycd* expression and high C3DC, whereas the opposite relationship was observed for C57BL/6J. (C) Quantification of malonic acid in urine samples from 129S2/SvPasCrl (n = 3 of which one pooled sample), DBA/2J (n = 2 pooled samples) and C57BL/6J (n = 4 of which one pooled sample) mice. (D) DNA gel electrophoresis after PCR amplification of a genomic region of the *Mlycd* gene using a forward primer in intron 2 and a reverse primer in exon 3. The left gel shows the PCR products using DNA isolated from C57BL/6J (B), DBA/2J (D) and 129S2/SvPasCrl (129). The right gel shows the PCR products using DNA isolated from C57BL/6J (B), DBA/2J (D) and SM/J (S). Bands are shown as the inverted colors of the ethidium bromide staining. The sequence of the DBA/2J fragment is available at GenBank under accession number MH036232. (E) Correlation between *Mlycd* expression in liver and the plasma levels of the ketone body derived C4OH-carnitine in samples from BXD mice on CD and HFD. (F) Schematic representation of malonyl-CoA metabolism and its interaction with fatty acid oxidation and ketogenesis.

**Figure S2. Weighted correlation network analysis of metabolites.** (A) WGCNA identified 5 modules of highly correlated metabolites; long-chain acylcarnitines (LCAC, turquoise), lipid (yellow), total amino acids (total AA, brown), branched-chain amino acids (BCAA, green) and short-chain acylcarnitines (SCAC, blue). Heatmaps showing the values of the metabolites in each module for each individual BXD strain under CD and HFD condition. (B) Individual eigenmetabolite values for each BXD strain under CD and HFD conditions and for each metabolite module. (C) The mean values (+ SEM) of the eigenmetabolites under CD and HFD condition for each module. (D) The correlation between the eigenmetabolites under CD and HFD condition for each module.

**Figure S3. Weighted correlation network analysis of liver and muscle gene expression.** (A) WGCNA of liver gene expression data revealed 20 modules of highly correlated genes in the CD condition (median size 120.5 genes) and revealed 17 modules of highly correlated genes in the

HFD condition (median size 100 genes). (B) WGCNA of muscle gene expression data revealed 31 modules of highly correlated genes in the CD condition (median size 74 genes) and revealed 17 modules of highly correlated genes in the HFD condition (median size 112 genes). Each matrix shows the size of each module on CD or HFD and the number of genes shared between CD and HFD modules with the associated FDR value. Only data for significant enrichments (FDR < 0.05) are shown. The eigengene value (PC1) was used to determine QTLs for each module. Significance of the QTL peak(s) are indicated $p < 0.1$ (*) or $p < 0.05$ (**). See also Table S18.

**Figure S4. Bayesian gene regulatory network analysis reveals glucocorticoid signaling as a candidate modifying pathway in FAO-deficient muscle.** (A) The normalized path length of the DEGs of the LCAD KO muscle as well as the genes of the MC4 and MH2 modules within the human and mouse muscle GRNs. See also Table S22. (B) Human subnetworks generated from LCAD KO muscle DEGs and the MC4 and MH2 modules share many genes (top) and are highly enriched in the original LCAD KO muscle DEGs (bottom; FE = fold enrichment).

**Figure S5. Identifying inflammatory pathways associated with phenotype variability in Gaucher disease.** The normalized path length of GD liver DEGs as well as the genes of the LC4, LH3, LH2 and LH4 modules within the human and mouse liver GRNs. See also Table S33.

**Supplementary Tables**

Rows with data important for the content of this manuscript are highlighted with a yellow fill color.

*File: supplemental_data_tables_1_to_9.xlsx*

Supplementary Table 1. Metabolite abundance values and annotations

Supplementary Table 2. Plasma and liver metabolite QTLs

Supplementary Table 3. Plasma and liver metabolite QTL (mQTL) lead SNPs

Supplementary Table 4. BXD gene expression QTLs (eQTLs): Liver CD

Supplementary Table 5. BXD gene expression QTLs (eQTLs): Liver HFD

Supplementary Table 6. BXD gene expression QTLs (eQTLs): Muscle CD

Supplementary Table 7. BXD gene expression QTLs (eQTLs): Muscle HFD

Supplementary Table 8. eQTLs and gene-trait correlations co-mapping to mQTLs

Supplementary Table 9. Diet-conserved eQTLs with significant gene-trait correlations

*File: supplemental_data_tables_10_to_18.xlsx*

Supplementary Table 10. LCAD KO mouse DEG signatures: muscle and liver

Supplementary Table 11. BXD metabolite coexpression module eigenvector values

Supplementary Table 12. BXD gene coexpression modules: liver and muscle, CD and HFD

Supplementary Table 13. BXD gene coexpression module conservation: liver CD and liver HFD; muscle CD and muscle HFD

Supplementary Table 14. BXD liver CD gene coexpression module eigenvector values

Supplementary Table 15. BXD liver HFD gene coexpression module eigenvector values

Supplementary Table 16. BXD muscle CD gene coexpression module eigenvector values

Supplementary Table 17. BXD muscle HFD gene coexpression module eigenvector values

Supplementary Table 18. BXD gene coexpression module QTLs

*File: supplemental_data_tables_19_to_21.xlsx*

Supplementary Table 19. BXD gene coexpression module enrichment with disease signatures: LCAD KO liver and muscle, mutant *Gba* liver

Supplementary Table 20. BXD gene coexpression module correlations with metabolite modules, liver metabolites, and clinical traits.

Supplementary Table 21. Gene set annotations of LCAD KO-associated modules with GO, KEGG, and Reactome gene sets

*File: supplemental_data_tables_22_to_31.xlsx*

Supplementary Table 22. Shortest path analysis for LCAD KO-related muscle signatures and modules

Supplementary Table 23. LCAD KO-specific subnetworks in muscle and liver

Supplementary Table 24. LCAD KO muscle subnetwork enrichment in LCAD KO muscle DEG signature

Supplementary Table 25. Overlapping muscle LCAD KO subnetwork genes

Supplementary Table 26. Annotating LCAD KO-related muscle subnetworks with enriched transcriptional regulatory motifs

Supplementary Table 27: LCAD KO key driver analysis in muscle and liver

Supplementary Table 28. Set of GR target genes

Supplementary Table 29. LCAD KO-related muscle signatures and subnetworks enriched in GR target genes

Supplementary Table 30. LCAD KO subnetworks enrichment in Klf15 KO muscle DEG signature

Supplementary Table 31. Statistical analysis of experimental data: GR signaling modulates response to fasting in pharmacological FAO disorder model

*File: supplemental_data_tables_32_to_42.xlsx*

Supplementary Table 32. Liver DEG signature from a Gba1 KO mouse model of Gaucher disease (Dasgupta, N et al, PLoS One, 2013, Liver_DESeq_9V/null saline vs. WT)

Supplementary Table 33. Shortest path analysis for Gaucher disease-related signatures and modules

Supplementary Table 34. Gaucher disease-specific liver subnetworks

Supplementary Table 35. Overlapping genes in Gaucher-specific subnetworks

Supplementary Table 36. Gaucher disease-specific liver subnetwork enrichment in Gaucher disease molecular signature

Supplementary Table 37. Gaucher disease-specific liver signatures, BXD modules, and subnetworks annotated with associated gene sets from WikiPathways and C5a signatures from GD patient-derived macrophages

Supplementary Table 38. Gaucher disease-specific liver signatures, BXD modules, and subnetworks annotated with canonical Hallmark pathways

Supplementary Table 39. Annotating Gaucher disease-specific liver subnetworks with enriched transcriptional regulatory motifs

Supplementary Table 40. Gaucher disease-specific liver signatures, BXD modules, and subnetworks annotated with canonical pathways in the Comparative Toxicogenomics Database (CTD)

Supplementary Table 41. Gaucher disease-specific key driver analysis

Supplementary Table 42. Cross-tissue correlation of BXD gene coexpression modules: liver and muscle

**Supplementary References**

[1] E.G. Williams, Y. Wu, P. Jha, S. Dubuis, P. Blattmann, C.A. Argmann, S.M. Houten, T. Amariuta, W. Wolski, N. Zamboni, R. Aebersold, J. Auwerx, Systems proteomics of liver mitochondria function, Science 352(6291) (2016) aad0189.
[2] Y. Wu, E.G. Williams, S. Dubuis, A. Mottis, V. Jovaisaite, S.M. Houten, C.A. Argmann, P. Faridi, W. Wolski, Z. Kutalik, N. Zamboni, J. Auwerx, R. Aebersold, Multilayered genetic and omics dissection of mitochondrial activity in a mouse reference population, Cell 158(6) (2014) 1415-30.
[3] C.K. Chuang, T.J. Wang, C.Y. Yeung, D.S. Lin, H.Y. Lin, H.L. Liu, H.T. Ho, W.S. Hsieh, S.P. Lin, A method for lactate and pyruvate determination in filter-paper dried blood spots, J Chromatogr A 1216(51) (2009) 8947-52.
[4] B. Casetta, D. Tagliacozzi, B. Shushan, G. Federici, Development of a method for rapid quantitation of amino acids by liquid chromatography-tandem mass spectrometry (LC-MSMS) in plasma, Clin Chem Lab Med 38(5) (2000) 391-401.
[5] M. Piraud, C. Vianey-Saban, K. Petritis, C. Elfakir, J.P. Steghens, A. Morla, D. Bouchu, ESI-MS/MS analysis of underivatised amino acids: a new tool for the diagnosis of inherited disorders of amino acid metabolism. Fragmentation study of 79 molecules of biological interest in positive and negative ionisation mode, Rapid Commun. Mass Spectrom. 17(12) (2003) 1297-1311.
[6] P. Vreken, A.E. van Lint, A.H. Bootsma, H. Overmars, R.J. Wanders, A.H. van Gennip, Quantitative plasma acylcarnitine analysis using electrospray tandem mass spectrometry for the diagnosis of organic acidaemias and fatty acid oxidation defects, J.Inherit.Metab Dis. 22(3) (1999) 302-306.
[7] J.E. Groener, B.J. Poorthuis, S. Kuiper, M.T. Helmond, C.E. Hollak, J.M. Aerts, HPLC for simultaneous quantification of total ceramide, glucosylceramide, and ceramide trihexoside concentrations in plasma, Clin Chem 53(4) (2007) 742-7.
[8] M. Mirzaian, P. Wisse, M.J. Ferraz, A.R.A. Marques, T.L. Gabriel, C. van Roomen, R. Ottenhoff, M. van Eijk, J.D.C. Codee, G.A. van der Marel, H.S. Overkleeft, J.M. Aerts, Accurate quantification of sphingosine-1-phosphate in normal and Fabry disease plasma, cells and tissues by LC-MS/MS with (13)C-encoded natural S1P as internal standard, Clinica chimica acta; international journal of clinical chemistry 459 (2016) 36-44.
[9] H. Gold, M. Mirzaian, N. Dekker, M. Joao Ferraz, J. Lugtenburg, J.D. Codee, G.A. van der Marel, H.S. Overkleeft, G.E. Linthorst, J.E. Groener, J.M. Aerts, B.J. Poorthuis, Quantification of globotriaosylsphingosine in plasma and urine of fabry patients by stable isotope ultraperformance liquid chromatography-tandem mass spectrometry, Clin Chem 59(3) (2013) 547-56.
[10] K.W. Broman, H. Wu, S. Sen, G.A. Churchill, R/qtl: QTL mapping in experimental crosses, Bioinformatics 19(7) (2003) 889-90.
[11] J. Leandro, A. Bender, T. Dodatko, C. Argmann, C. Yu, S.M. Houten, Glutaric aciduria type 3 is a naturally occurring biochemical trait in inbred mice of 129 substrains, Mol Genet Metab 132(2) (2021) 139-145.
[12] J. Leandro, S. Violante, C.A. Argmann, J. Hagen, T. Dodatko, A. Bender, W. Zhang, E.G. Williams, A.M. Bachmann, J. Auwerx, C. Yu, S.M. Houten, Mild inborn errors of metabolism in commonly used inbred mouse strains, Mol Genet Metab 126(4) (2019) 388-396.

[13] A. Dobin, C.A. Davis, F. Schlesinger, J. Drenkow, C. Zaleski, S. Jha, P. Batut, M. Chaisson, T.R. Gingeras, STAR: ultrafast universal RNA-seq aligner, Bioinformatics 29(1) (2013) 15-21.
[14] Y. Liao, G.K. Smyth, W. Shi, featureCounts: an efficient general purpose program for assigning sequence reads to genomic features, Bioinformatics 30(7) (2014) 923-30.
[15] J.T. Robinson, H. Thorvaldsdottir, W. Winckler, M. Guttman, E.S. Lander, G. Getz, J.P. Mesirov, Integrative genomics viewer, Nat Biotechnol 29(1) (2011) 24-6.
[16] H. Thorvaldsdottir, J.T. Robinson, J.P. Mesirov, Integrative Genomics Viewer (IGV): high-performance genomics data visualization and exploration, Brief Bioinform 14(2) (2013) 178-92.
[17] P. Ranea-Robles, C. Yu, N. van Vlies, F.M. Vaz, S.M. Houten, Slc22a5 haploinsufficiency does not aggravate the phenotype of the long-chain acyl-CoA dehydrogenase KO mouse, J Inherit Metab Dis 43(3) (2020) 486-495.
[18] N. van Vlies, L. Tian, H. Overmars, A.H. Bootsma, W. Kulik, R.J. Wanders, P.A. Wood, F.M. Vaz, Characterization of carnitine and fatty acid metabolism in the long-chain acyl-CoA dehydrogenase-deficient mouse, Biochem. J. 387(Pt 1) (2005) 185-193.
[19] E.F. Diekman, M. van Weeghel, R.J. Wanders, G. Visser, S.M. Houten, Food withdrawal lowers energy expenditure and induces inactivity in long-chain fatty acid oxidation-deficient mouse models, FASEB J. 28(7) (2014) 2891-2900.
[20] M.I. Love, W. Huber, S. Anders, Moderated estimation of fold change and dispersion for RNA-seq data with DESeq2, Genome Biol 15(12) (2014) 550.
[21] C.A. Argmann, S. Violante, T. Dodatko, M.P. Amaro, J. Hagen, V.L. Gillespie, C. Buettner, E.E. Schadt, S.M. Houten, Germline deletion of Kruppel-like factor 14 does not increase risk of diet induced metabolic syndrome in male C57BL/6 mice, Biochim Biophys Acta (2017).
[22] N. Dasgupta, Y.H. Xu, S. Oh, Y. Sun, L. Jia, M. Keddache, G.A. Grabowski, Gaucher disease: transcriptome analyses using microarray or mRNA sequencing in a Gba1 mutant mouse model treated with velaglucerase alfa or imiglucerase, PloS one 8(10) (2013) e74912.
[23] B. Zhang, S. Horvath, A general framework for weighted gene co-expression network analysis, Statistical applications in genetics and molecular biology 4 (2005) Article17.
[24] J.A. Miller, C. Cai, P. Langfelder, D.H. Geschwind, S.M. Kurian, D.R. Salomon, S. Horvath, Strategies for aggregating gene expression data: the collapseRows R function, BMC bioinformatics 12 (2011) 322.
[25] E.E. Schadt, C. Molony, E. Chudin, K. Hao, X. Yang, P.Y. Lum, A. Kasarskis, B. Zhang, S. Wang, C. Suver, J. Zhu, J. Millstein, S. Sieberts, J. Lamb, D. GuhaThakurta, J. Derry, J.D. Storey, I. Avila-Campillo, M.J. Kruger, J.M. Johnson, C.A. Rohl, N.A. van, M. Mehrabian, T.A. Drake, A.J. Lusis, R.C. Smith, F.P. Guengerich, S.C. Strom, E. Schuetz, T.H. Rushmore, R. Ulrich, Mapping the genetic architecture of gene expression in human liver, PLoS.Biol. 6(5) (2008) e107.
[26] J. Zhu, P.Y. Lum, J. Lamb, D. GuhaThakurta, S.W. Edwards, R. Thieringer, J.P. Berger, M.S. Wu, J. Thompson, A.B. Sachs, E.E. Schadt, An integrative genomics approach to the reconstruction of gene networks in segregating populations, Cytogenetic and genome research 105(2-4) (2004) 363-74.

[27] J. Zhu, M.C. Wiener, C. Zhang, A. Fridman, E. Minch, P.Y. Lum, J.R. Sachs, E.E. Schadt, Increasing the power to detect causal associations by combining genotypic and expression data in segregating populations, PLoS.Comput.Biol. 3(4) (2007) e69.
[28] J. Zhu, P. Sova, Q. Xu, K.M. Dombek, E.Y. Xu, H. Vu, Z. Tu, R.B. Brem, R.E. Bumgarner, E.E. Schadt, Stitching together multiple data dimensions reveals interacting metabolomic and transcriptomic networks that modulate cell regulation, PLoS biology 10(4) (2012) e1001301.
[29] S. Durinck, Y. Moreau, A. Kasprzyk, S. Davis, B. De Moor, A. Brazma, W. Huber, BioMart and Bioconductor: a powerful link between biological databases and microarray data analysis, Bioinformatics 21(16) (2005) 3439-40.
[30] H. Attrill, P. Gaudet, R.P. Huntley, R.C. Lovering, S.R. Engel, S. Poux, K.M. Van Auken, G. Georghiou, M.C. Chibucos, T.Z. Berardini, V. Wood, H. Drabkin, P. Fey, P. Garmiri, M.A. Harris, T. Sawford, L. Reiser, R. Tauber, S. Toro, C. Gene Ontology, Annotation of gene product function from high-throughput studies using the Gene Ontology, Database (Oxford) 2019 (2019).
[31] M. Kanehisa, S. Goto, KEGG: kyoto encyclopedia of genes and genomes, Nucleic Acids Res 28(1) (2000) 27-30.
[32] A. Fabregat, K. Sidiropoulos, G. Viteri, O. Forner, P. Marin-Garcia, V. Arnau, P. D'Eustachio, L. Stein, H. Hermjakob, Reactome pathway analysis: a high-performance in-memory approach, BMC bioinformatics 18(1) (2017) 142.
[33] G. Bindea, B. Mlecnik, H. Hackl, P. Charoentong, M. Tosolini, A. Kirilovsky, W.H. Fridman, F. Pages, Z. Trajanoski, J. Galon, ClueGO: a Cytoscape plug-in to decipher functionally grouped gene ontology and pathway annotation networks, Bioinformatics 25(8) (2009) 1091-3.
[34] A. Liberzon, C. Birger, H. Thorvaldsdottir, M. Ghandi, J.P. Mesirov, P. Tamayo, The Molecular Signatures Database (MSigDB) hallmark gene set collection, Cell Syst 1(6) (2015) 417-425.
[35] C.J. Mattingly, M.C. Rosenstein, G.T. Colby, J.N. Forrest, Jr., J.L. Boyer, The Comparative Toxicogenomics Database (CTD): a resource for comparative toxicological studies, J Exp Zool A Comp Exp Biol 305(9) (2006) 689-92.
[36] T. Kuo, M.J. Lew, O. Mayba, C.A. Harris, T.P. Speed, J.C. Wang, Genome-wide analysis of glucocorticoid receptor-binding sites in myotubes identifies gene networks modulating insulin signaling, Proc Natl Acad Sci USA 109(28) (2012) 11160-5.
[37] R. Janky, A. Verfaillie, H. Imrichova, B. Van de Sande, L. Standaert, V. Christiaens, G. Hulselmans, K. Herten, M. Naval Sanchez, D. Potier, D. Svetlichnyy, Z. Kalender Atak, M. Fiers, J.C. Marine, S. Aerts, iRegulon: from a gene list to a gene regulatory network using large motif and track collections, PLoS computational biology 10(7) (2014) e1003731.
[38] S. Gray, B. Wang, Y. Orihuela, E.G. Hong, S. Fisch, S. Haldar, G.W. Cline, J.K. Kim, O.D. Peroni, B.B. Kahn, M.K. Jain, Regulation of gluconeogenesis by Kruppel-like factor 15, Cell Metab 5(4) (2007) 305-12.
[39] J.C. Serfecz, A. Saadin, C.P. Santiago, Y. Zhang, S.M. Bentzen, S.N. Vogel, R.A. Feldman, C5a Activates a Pro-Inflammatory Gene Expression Profile in Human Gaucher iPSC-Derived Macrophages, Int J Mol Sci 22(18) (2021).

[40] B. Zhang, J. Zhu, Identification of Key Causal Regulators in Gene Networks, in: S.I. Ao, L. Gelman, D.W.L. Hukins, A. Hunter, A.M. Korsunsky (Eds.) The World Congress on Engineering, Newswood Limited, London, U.K., 2013, pp. 1309-1312.
[41] P. Ranea-Robles, S. Violante, C. Argmann, T. Dodatko, D. Bhattacharya, H. Chen, C. Yu, S.L. Friedman, M. Puchowicz, S.M. Houten, Murine deficiency of peroxisomal L-bifunctional protein (EHHADH) causes medium-chain 3-hydroxydicarboxylic aciduria and perturbs hepatic cholesterol homeostasis, Cellular and molecular life sciences : CMLS 78(14) (2021) 5631-5646.
[42] S. Violante, N. Achetib, C.W.T. van Roermund, J. Hagen, T. Dodatko, F.M. Vaz, H.R. Waterham, H. Chen, M. Baes, C. Yu, C.A. Argmann, S.M. Houten, Peroxisomes can oxidize medium- and long-chain fatty acids through a pathway involving ABCD3 and HSD17B4, FASEB J 33(3) (2019) 4355-4364.

### Supplementary Figure 1

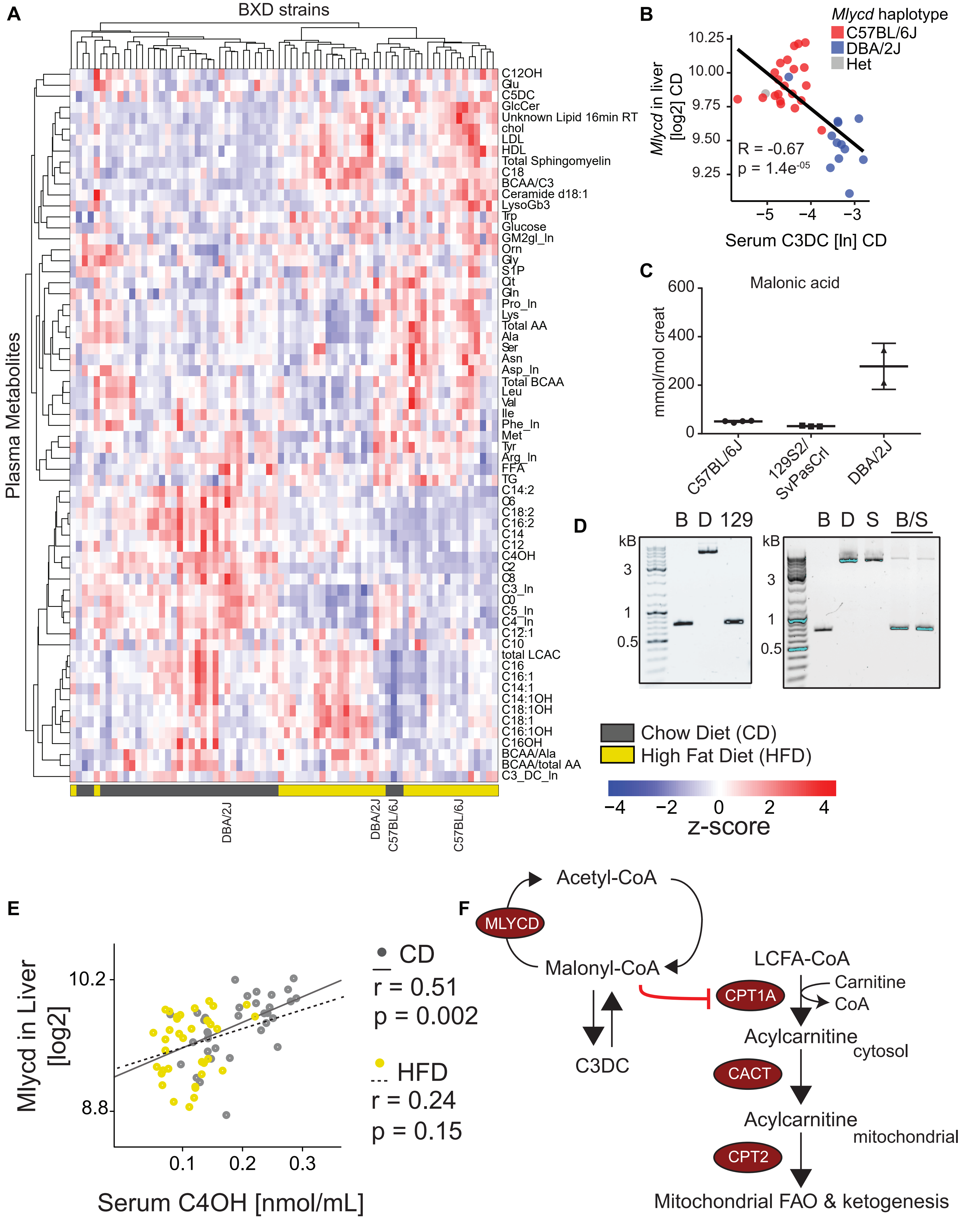

### Supplementary Figure 2

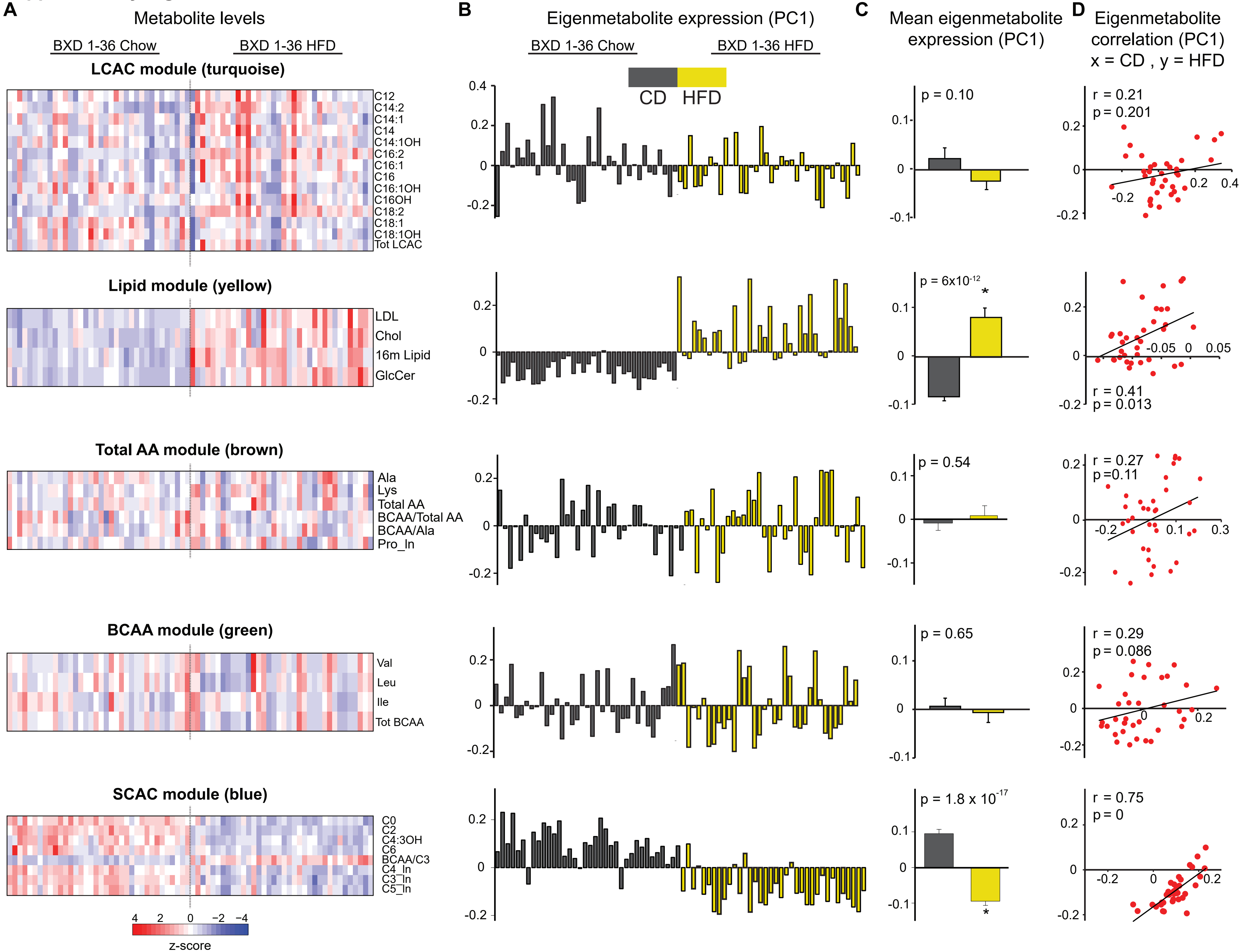

**Supplementary Figure 3**

**A.**

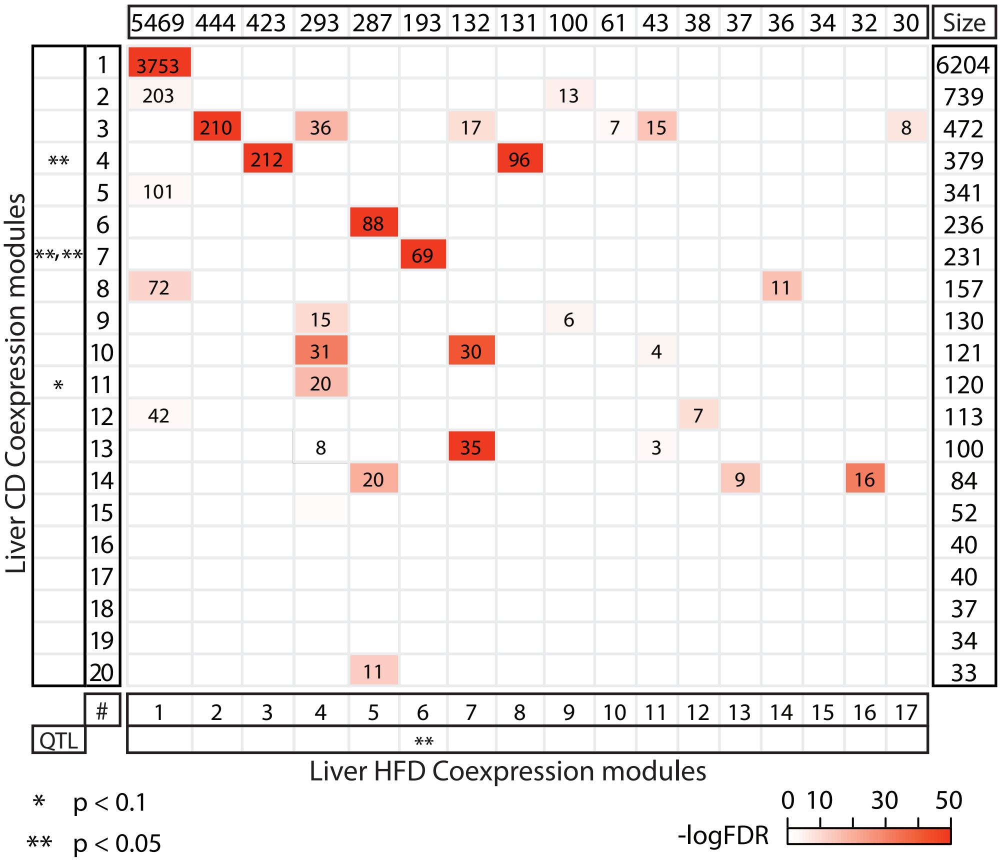


**B.**

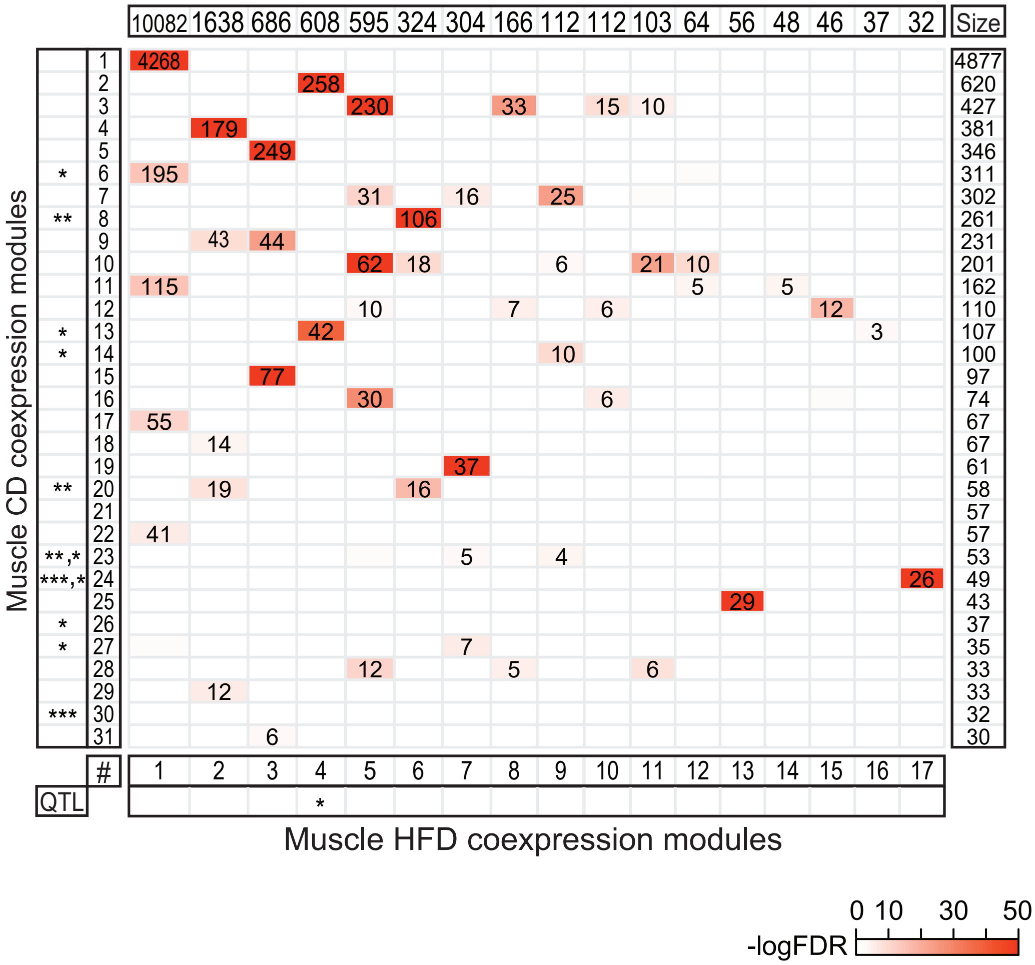

**Supplementary Figure 4**

A

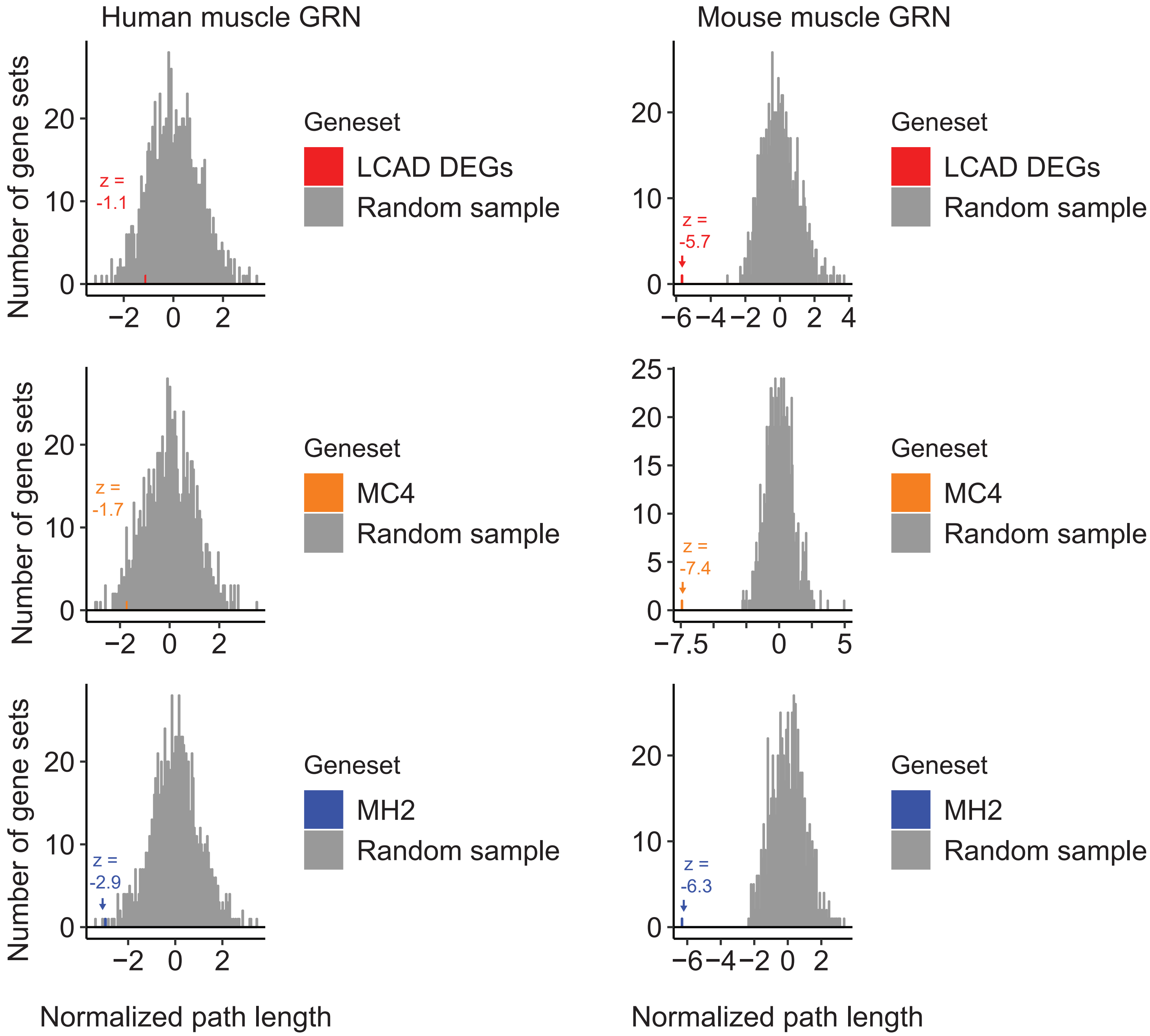


B

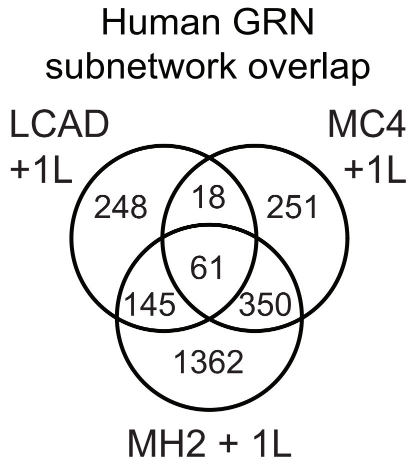

**Supplementary Figure 5**

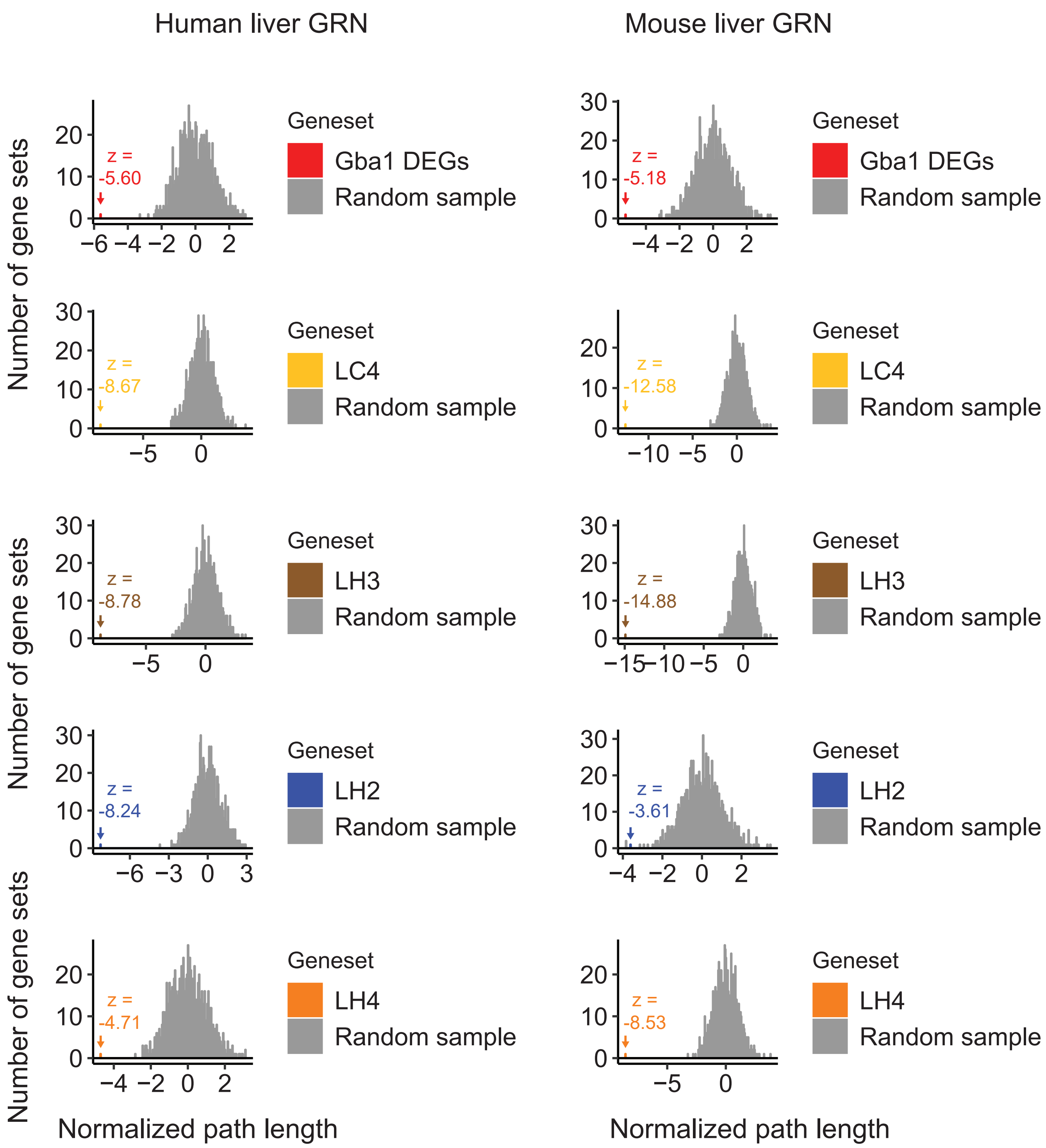
